# Transcriptional activation of auxin biosynthesis drives developmental reprogramming of differentiated cells

**DOI:** 10.1101/2021.06.26.450054

**Authors:** Yuki Sakamoto, Ayako Kawamura, Takamasa Suzuki, Shoji Segami, Masayoshi Maeshima, Stefanie Polyn, Lieven De Veylder, Keiko Sugimoto

## Abstract

Plant cells exhibit remarkable plasticity of their differentiation states, enabling regeneration of whole plants from differentiated somatic cells. How they revert cell fate and express pluripotency, however, remains unclear. Here we show that transcriptional activation of auxin biosynthesis is crucial for reprogramming differentiated Arabidopsis leaf cells. We demonstrate that interfering with the activity of histone acetyltransferases dramatically reduces callus formation from leaf mesophyll protoplasts. Impaired histone acetylation predominantly affects transcription of auxin biosynthesis genes. Auxin biosynthesis is in turn required to accomplish initial cell division through the activation of G2/M phase genes mediated by MYB DOMAIN PROTEIN 3-RELATED (MYB3Rs). We further show that the AUXIN RESPONSE FACTOR 7 (ARF7)/ARF19 and INDOLE-3-ACETIC ACID INDUCIBLE 3 (IAA3)/IAA18-mediated auxin signaling pathway is responsible for cell cycle reactivation in protoplasts. These findings provide novel mechanistic model of how differentiated plant cells can revert their fate and reinitiate the cell cycle to become pluripotent.

---

Tight coordination of cell proliferation and differentiation is central to optimize plant organ development. Typically, during a normal developmental program, meristematic cells continue to proliferate until they start to differentiate, and once differentiated, the somatic cells usually remain mitotically inactivate^1–3^. Such a relationship between mitosis and cellular differentiation status should be also critical during developmental reprogramming through which plant somatic cells convert their fates and regenerate new tissues or organs. Previous studies indeed suggest that reactivation of somatic cell division contributes to cell fate conversion and acquisition of pluripotency presumably by helping cells dilute existing identity and activate a new developmental program^4–6^. Uncovering how differentiated cells reinitiate cell division is therefore crucial to understand how somatic cells initiate reprogramming.

Mesophyll cells in mature leaves have fully developed organelles such as chloroplasts, and they usually do not divide in intact plant tissues. When they are isolated by cell wall digestion as single cells called protoplasts and cultivated under phytohormone-containing conditions, however, they reinitiate the cell cycle, grow into an unorganized cell mass called callus and even regenerate whole plants^7^. It is thus clearly demonstrated that differentiated leaf cells can change their fate and take on a meristematic, pluripotent state. Previous studies have described the cytological and physiological properties of protoplasts during cell cycle reinitiation in several plant species, including tobacco (*Nicotiana tabacum*) and Arabidopsis (*Arabidopsis thaliana*). Protoplasts, for instance, undergo cell wall reconstruction^8,9^ and changes in the structure and subcellular localization of organelles^10,11^. At the physiological level, cell cycle reinitiation is known to correlate with the neutralization of reactive oxygen species (ROS)^12,13^ and phytohormone production^14^. Additionally, several studies have demonstrated that freshly isolated protoplasts possess a more opened chromatin state compared to intact leaf cells, possibly caused by drastic changes in epigenetic modifications^15,16^. Consistently, Chupeau *et al.*^17^ reported that Arabidopsis protoplasts undergo dynamic transcriptomic reprogramming during early phases of incubation. Despite these reported findings, the molecular mechanisms that drive cell cycle reinitiation from differentiated cells remain obscure, partly due to a lack of experimental systems to quantitatively assess this cellular process. To overcome this problem, we improved the culture system for Arabidopsis leaf mesophyll protoplasts and then performed quantitative genetic, physiological and cell biological analyses. Using these approaches, we demonstrate that transcriptional activation of auxin biosynthesis is essential to promote the cell cycle reactivation in differentiated cells.

## Results

### Arabidopsis leaf mesophyll protoplasts reprogram and regenerate shoots *in vitro*

We first established a new experimental pipeline to reproducibly induce callus formation from Arabidopsis leaf protoplasts (Fig. 1a). Protoplasts were isolated from mature rosette leaves with high viability, embedded into sodium alginate gels and cultured in protoplast callus induction medium (PCIM) supplemented with 2,4-D and thidiazuron as auxin and cytokinin, respectively (Fig. 1a, Supplementary Fig. 1a and Supplementary Table 1). After a 14-day incubation, around 2% of total embedded protoplasts develop into callus, enabling quantitative evaluation of this phenotype (Supplementary Fig.1b). When these protoplast-derived calli are transferred to callus growth medium (CGM) and subsequently to shoot induction medium (SIM), they continue to proliferate and regenerate shoots (Fig. 1b), indicating that these protoplasts acquire competence to form shoot meristems.

**Figure 1.**
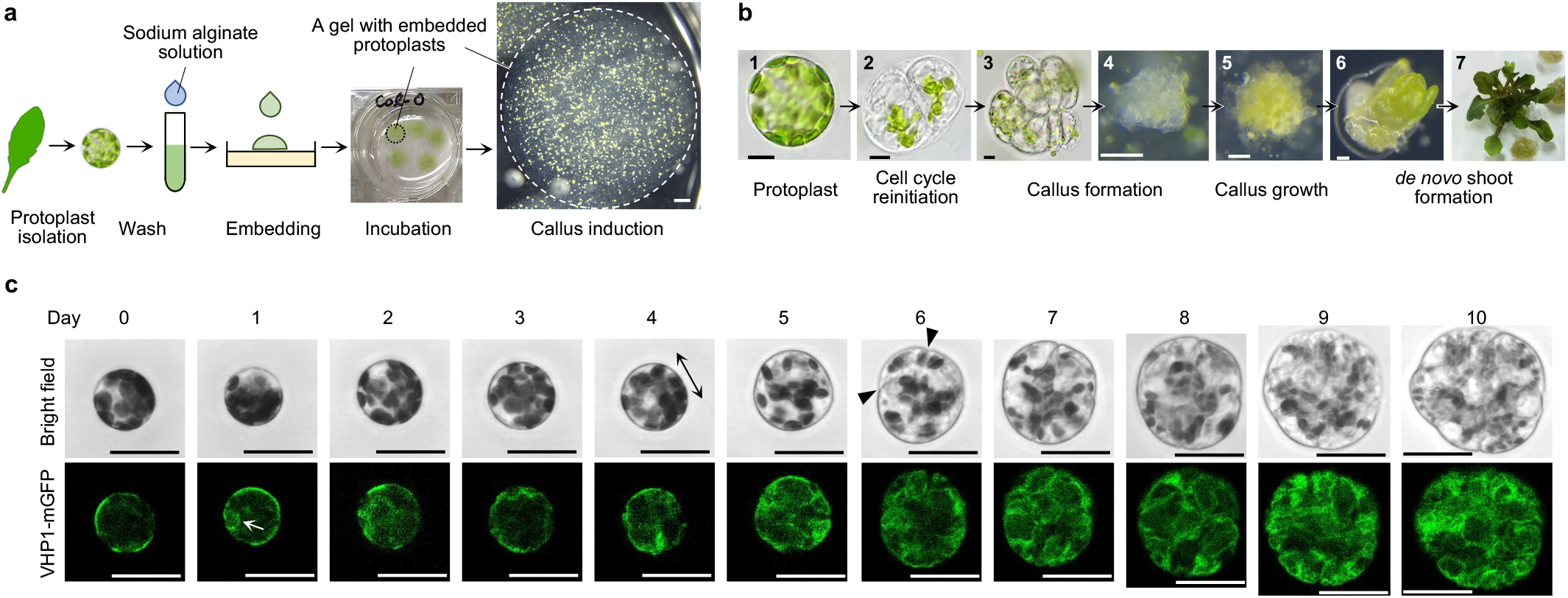
Leaf mesophyll protoplasts reprogram into a pluripotent state and regenerate shoots *in vitro*. **a**, Diagram showing the key steps for protoplast isolation and callus induction. The circles with dashed lines indicate a sodium alginate gel that contains protoplasts. **b**, Light microscopy images of a freshly isolated leaf mesophyll protoplast (**1**), a protoplast that has undergone the first cell division (**2**), callus formed from a protoplast (**3** to **5**), and new shoots formed from protoplast-derived callus (**6**, **7**). **c**, Time-lapse confocal microscopy images of a leaf mesophyll protoplast undergoing cellular reprogramming from Day 0 to Day 10. Vacuolar morphology is visualized by VHP1-mGFP. The double-headed arrow indicates the direction of cell elongation and arrowheads mark the plane of initial cell division. The white arrow highlights the initial appearance of vacuolar strand-like structures. Scale bars are 10 μm (**1** to **3** in **a**), 100 μm (**4** to **6** in **a**), 1 mm (**b**) and 30 μm (**c**).

To uncover the cellular basis for developmental reprogramming, we first determined the original identity of dividing protoplasts. The *pCAB3:H2A-eGFP* reporter, which drives nuclear-localized accumulation of HISTONE 2A-6 (H2A) fused with enhanced green fluorescent protein (eGFP) under the control of a promoter sequence of a mesophyll cell-specific gene *CHLOROPHYLL A/B BINDING PROTEIN 3* (*CAB3*)^18^, is expressed in most freshly isolated protoplasts (Supplementary Fig. 1c). Furthermore, using a time-lapse live imaging system that tracks reprogramming of individual protoplasts over time (Supplementary Fig. 1d), we confirmed that 98.78% (162 out of 164) of protoplasts that undergo cell division display mesophyll cell-like characteristics at Day 0 based on H2A-eGFP expression and/or the appearance and density of chloroplasts^19^ (Supplementary Fig. 1e-g). The two remaining H2A-eGFP-negative protoplasts also contain chloroplasts but they resemble guard cells based on chloroplast density^19^ (Supplementary Fig. 1h). These observations establish that the majority of protoplasts in our experimental setup initially have mesophyll cell identities.

Closer examination of early morphological changes via time-lapse imaging revealed that protoplasts undergo their first cell division after Day 4, and consistent with Chupeau *et al.*^17^, many divide at Day 6 or Day 7 (Fig1c, Supplementary Fig. 1i). As described previously^8,9^, most of them elongate anisotropically for 1 to 3 days before initial cell division (Supplementary Fig. 1j-k). We also observed protoplasts that elongate or expand isotopically without cell division as well as those that shrink or show no apparent changes in size or shape (Supplementary Fig. 2a-b). We have previously shown that vacuolar morphology, visualized by monomeric GFP (mGFP)-tagged VACUOLAR H^+^-PYROPHOSPHATASE 1 (VHP1), changes dynamically as cells transit from proliferative to differentiated phases^20^. Time-lapse imaging of protoplasts carrying *pVHP1:VHP1-mGFP* showed that freshly isolated protoplasts have a single large vacuole occupying most of the cell volume^11^ (Day 0 in Fig. 1c and Supplementary Fig. 2c). Strand-like structures, however, start to appear inside the vacuole either before or when the cells start to elongate and vacuoles undergo extensive compartmentalization as cells progress through successive divisions (Fig. 1c and Supplementary Fig. 2c). We observed similar strand-like structures in protoplasts that elongate or expand without cell division but they do not undergo or maintain similar levels of compartmentalization (Supplementary Fig. 2d-e), suggesting that the sustained vacuolar compartmentalization marks protoplasts reprogrammed to divide.

In parallel with the reactivation of cell proliferation, cellular dedifferentiation, i.e. the loss of existing traits, is another important aspect of cellular reprogramming. It is previously reported that during *in vitro* transdifferentiation of mesophyll cells into xylem cells, the expression levels of genes that characterize mesophyll cells, e.g. photosynthetic genes such as *CAB3* and *RIBULOSE BISPHOSPHATE CARBOXYLASE SMALL CHAIN 1A* (*RBCS1A*), are immediately decreased, reflecting the loss of mesophyll identity^21^. To investigate when protoplasts start to dedifferentiate, we examined the expression of genes involved in chloroplast functions in mature leaves and protoplasts at early incubation steps by RNA sequencing (Supplementary Fig.3 and Supplementary Table 2). We found that many genes encoding photosynthetic components, including subunits of light harvesting complexes, exhibit striking downregulation during protoplast isolation and/or following several days of incubation. In contrast, genes involved in chloroplast fission, the step important for chloroplast inheritance during cell division, are upregulated within 2 days of incubation, implying that protoplasts initiate transcription to prepare for cell cycle reinitiaton by this time. Importantly, many chloroplast-related genes that become downregulated in cultured protoplasts are known to be significantly upregulated as cells transit from proliferative to differentiated phases in growing Arabidopsis leaves^2^ (Supplementary Fig. 3), implying that the changes in expression of these genes in protoplasts reflects initiation of cellular dedifferentiation. These results thus suggest that cellular dedifferentiation starts as early as during protoplast isolation and the early steps of culture prior to initial cell division.

### Histone acetylation is required for cell cycle reinitiation in protoplasts

To elucidate the molecular mechanisms of developmental reprogramming, we next focused on the epigenetic modifications that protoplasts undergo during the process. Williams *et al.*^16^ reported that levels of histone acetylation, which is thought to promote gene expression^22^, increase in freshly isolated protoplasts compared to intact leaves. This suggests that histone acetylation ushers in transcriptional changes in protoplasts during early stages of culture, thus driving developmental reprogramming. Given that GNAT/MYST family histone acetyltransferases (HATs) regulate the expression of key genes in several regeneration contexts, such as during wound-induced callus formation and *in vitro* shoot regeneration from explants^23,24^, we first tested whether an inhibitor of this family of HATs, MB-3, interferes with protoplast reprograming. As shown in Fig. 2a-b, application of MB-3 to wild-type (WT) protoplasts strongly reduces callus formation efficiency at Day 14, with protoplasts largely failing to reinitiate cell division. Applying MB-3 at later time points causes similar defects in callus formation, suggesting that histone acetylation is also required for successive cell divisions (Supplementary Fig. 4a). Additionally, an inhibitor of CBP-family HATs, C646, strongly prevents callus formation, whereas garcinol, an inhibitor for p300 and PCAF HATs in human, has a much milder effect (Supplementary Fig. 4b). Furthermore, we found that among 12 HATs in Arabidopsis, HISTONE ACETYLTRANSFERASE OF THE GNAT/MYST SUPERFAMILY 1 (HAG1), HAG3 and HISTONE ACETYLATION OF THE TAF□250 FAMILY 1 (HAF1) are the key HATs involved in protoplast reprogramming since their single mutants are severely impaired in callus formation (Fig 2a, Supplementary Fig. 4c-e).

**Figure 2.**
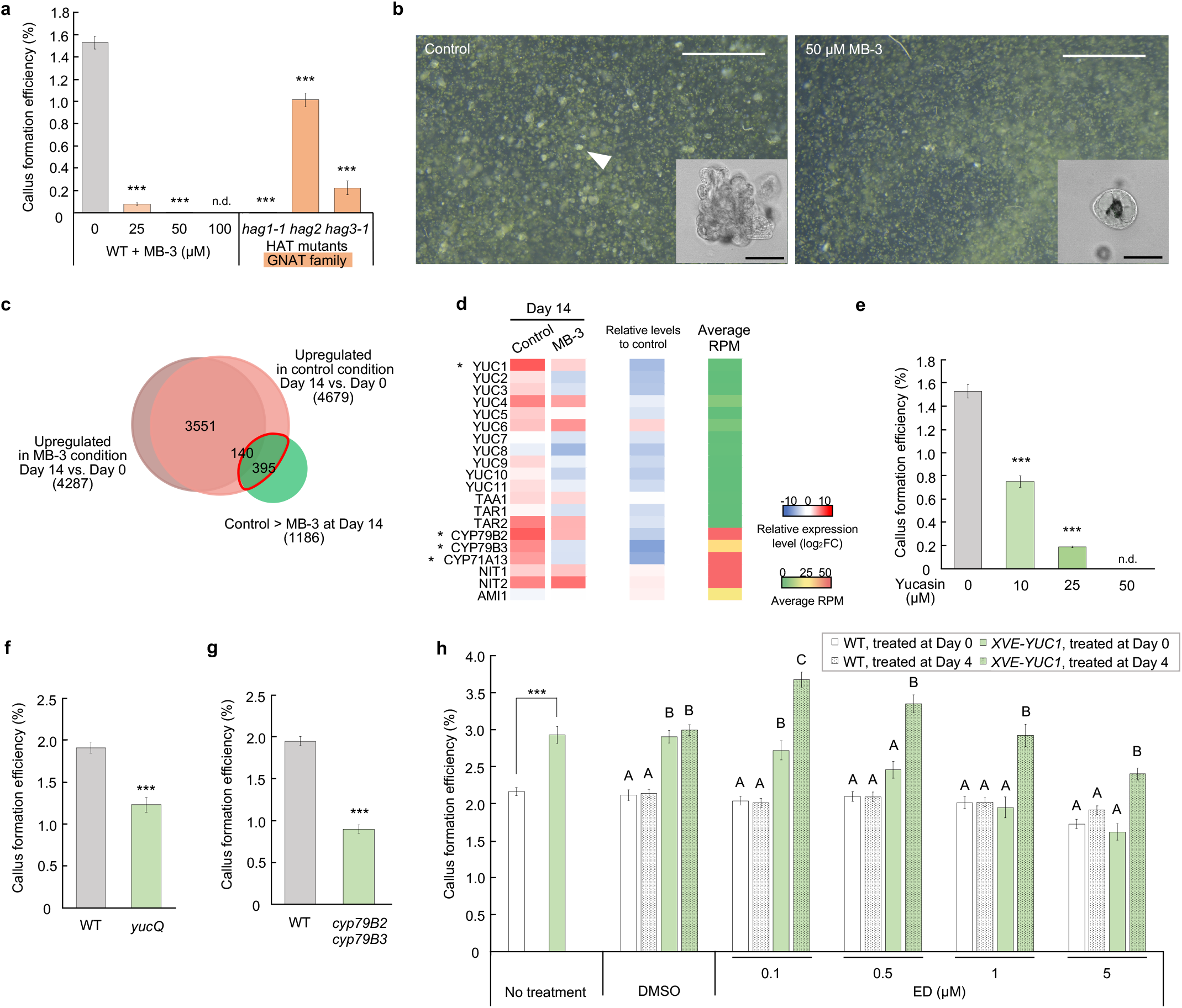
Histone acetylation is required to reinitiate cell division in leaf mesophyll protoplasts. **a**, Callus formation efficiency of MB3-treated WT, *hag1-1*, *hag2* and *hag3-1* protoplasts at Day 14. MB-3 was added at the beginning of culture (Day 0). Error bars represent standard error. n = 40 from 8 biological replicates for the WT control, 15 from 3 biological replicates for *hag1-1* and *hag2*, and 20 from 4 biological replicates for all others. \*\*\**P* < 0.001 (two-tailed Welch’s *t*-test compared to WT control). n.d. not determined. **b**, Bright field images of WT protoplasts incubated in control (left panel) and 50 μM MB-3 condition (right panel) for 14 days. The white arrowhead indicates a single callus. The bottom right insets show confocal microscopy images of callus cells (left panel) and an undivided protoplast (right panel). **c**, A Venn diagram of genes significantly upregulated in control and 50 μM MB-3 conditions. The green circle indicates genes that show significantly higher expression in the control compared to the MB-3 condition at Day 14. Numbers in brackets show total numbers of genes included in each group. The red outline highlights 535 genes that show significantly stronger upregulation in the control compared to the MB-3 condition. **d**, Heat map of the transcriptional changes for representative genes implicated in IAA biosynthesis. The left two columns show the expression levels in the control and MB-3 conditions at Day 14 as values normalized (log_2_FC) to Day 0. The ‘Relative levels to control’ column shows the differential expression levels in the MB-3 condition compared to the control at Day 14 as relative values (log_2_FC). Average reads per million (RPM) indicates the overall expression level for each gene. Asterisks mark the genes that are included from the 535 upregulated genes in **c**. **e**, Callus formation efficiency of yucasin-treated WT protoplasts. Error bars represent standard error. n = 30 from 6 biological replicates for the WT control and 15 from 3 biological replicates for the others. \*\*\**P* < 0.001 (two-tailed Student’s *t*-test or Welch’s *t*-test compared to WT control). n.d. not determined. **f**, Callus formation efficiency of WT and *yuc3 yuc5 yuc7 yuc8 yuc9* (*yucQ*) protoplasts. Error bars represent standard error. n = 15 from 3 biological replicates. \*\*\**P* < 0.001 (two-tailed Student’s *t*-test compared to WT). **g**, Callus formation efficiency of WT and *cyp79B2 cyp79B3* protoplasts. Error bars represent standard error. n = 15 from 3 biological replicates. \*\*\**P* < 0.001 (two-tailed Student’s *t*-test compared to WT). **h,** Callus formation efficiency of *XVE-YUC1* protoplasts. DMSO or β-estradiol (ED) were added to PCIM at the indicated final concentration at the beginning (Day 0) or Day 4 of culture. Error bars represent standard error. n = 15 from 3 biological replicates. For ’No treatment’, \*\*\**P* < 0.001 (two-tailed Welch’s *t*-test compared to WT under the same treatment). For the others, different letters indicate significant differences based on one-way ANOVA with post hoc Tukey’s HSD test (*P* < 0.001) within each treatment group. Scale bars are 1 mm (**b**) and 50 μm (insets in **b**).

To investigate how histone acetylation regulates protoplast reprograming, we examined gene expression via RNA-seq in freshly isolated protoplasts as well as in those cultured with or without 50 µM MB-3. Considering that the cell division occurs in only a few percent of protoplasts (Supplementary Fig. 2a) and is not synchronized over time (Supplementary Fig. 1i-j), we harvested the protoplasts cultured for 14 days, when we expect that the majority of division-competent protoplasts have undergone the first cell division, to maximize the number of detectable differentially expressed genes. Since histone acetylation is associated with activation of transcription, we focused on the 535 genes that show significantly stronger upregulation in the control condition compared to the MB-3 condition (Fig. 2c and Supplementary Table 3). Gene ontology (GO) analysis of the 535 upregulated genes revealed strong fold enrichments of the genes implicated in biosynthesis or metabolism of indole-containing compounds, especially tryptophan-derived ones such as indole glucosinolate and auxin indole-3-acetic acid (IAA) (Supplementary Table 4 and 5). More detailed examination of the expression patterns for a comprehensive set of genes implicated in tryptophan metabolism revealed that some IAA biosynthesis genes, including *YUCCA1* (*YUC1*) and *CYTOCHROME P450* genes (*CYP*s), are expressed at higher levels in the control condition compared to the MB-3 condition (Fig. 2d and Supplementary Table 6a). Although statistically not significant, we also observed upregulation of more numbers of other *YUC*s in the control condition, suggesting that IAA biosynthesis is repressed by MB-3 treatment. To test if IAA biosynthesis is required for cell cycle reinitiation, we examined whether blocking synthesis of this hormone impedes callus formation. As shown in Fig. 2e and Supplementary Fig. 5a, callus formation is strongly compromised in WT protoplasts treated with the IAA biosynthesis inhibitors yucasin and Kyn. We also observed similar defects in two mutants defective in IAA biosynthesis, *yuc3 yuc5 yuc7 yuc8 yuc9* (*yucQ*)^25^ and *cytochrome p450 family 79 subfamily B polypeptide 2* (*cyp79B2*) *cyp79B3*^26^ (Fig. 2f-g), supporting the idea that IAA biosynthesis promotes protoplast division. Consistently, overexpression of *YUC1* in *LexA-VP16-estrogen receptor (XVE)-YUC1* protoplasts or application of low concentrations of IAA reproducibly increases callus formation efficiency (Fig. 2h and Supplementary Fig. 5b-c), further substantiating that the level of IAA is a key limiting factor for cell cycle reactivation. Interestingly, it seems that *YUC* expression within an appropriate dose range is needed to maximally promote cell cycle reinitiation. A low level of leaky *YUC1* expression under mock treatment or mild induction of the *YUC1* transgene by 0.1 μM β-estradiol (ED) in *XVE-YUC1* are sufficient to increase callus formation efficiency compared to the WT, while strong overexpression of the *YUC1* transgene by 1 or 5 μM ED does not have this effect (Fig. 2h and Supplementary Fig. 5c). Furthermore, our data suggest that the timing of *YUC* expression is also important since induction of *YUC1* transgene at Day 4 results in greater promotion of callus formation compared to its induction at Day 0 (Fig. 2h and Supplementary Fig. 5c).

### Auxin biosynthesis specifically regulates initial division of protoplasts by activating their auxin response

Our results so far suggest that IAA biosynthesis prior to the initial cell division promotes callus formation from mesophyll protoplasts. Indeed, our RNA-seq data at early time points show that IAA biosynthesis genes are upregulated before initial cell division (Fig. 3a and Supplementary Table 6b). These data support the previous findings that the IAA content is transiently increased just before initial cell division in alfalfa protoplasts^14^. We therefore hypothesized that IAA biosynthesis specifically promotes initial cell division. Notably, yucasin strongly inhibits callus formation only when it is added by Day 4, i.e. before most cells reinitiate the cell cycle (Fig. 3b), indicating that endogenous IAA production is specifically required for initial cell division. To further investigate the role of IAA biosynthesis during early stages of protoplast reprogramming, we visualized the auxin response by time-lapse imaging of protoplasts carrying *DR5rev:GFP*, which expresses GFP via the auxin response element-containing *DR5* promoter^27,28^. In the control condition, we barely observed *DR5rev:GFP* expression in freshly isolated protoplasts, while many start to express detectable GFP signals from Day 2 to Day 4 (Fig. 3c-d), with nearly 50% of protoplasts showing *DR5rev:GFP* expression at least at one time point by Day 6. Importantly, we further found that IAA biosynthesis is required for activation of the auxin response, since yucasin treatment severely reduces the proportion of *DR5rev:GFP*-positive protoplasts (Fig. 3d). We should also note that *DR5rev:GFP*-positive cells in the control condition are enriched among protoplasts that divide, elongate or expand, whereas *DR5rev:GFP*-negative protoplasts in either the control or yucasin condition generally decrease in size (Supplementary Fig. 6a-b). These observations suggest that *DR5rev:GFP*-detectable auxin response promotes the early phases of developmental reprograming although that alone is not sufficient to complete the first cell division.

**Figure 3.**
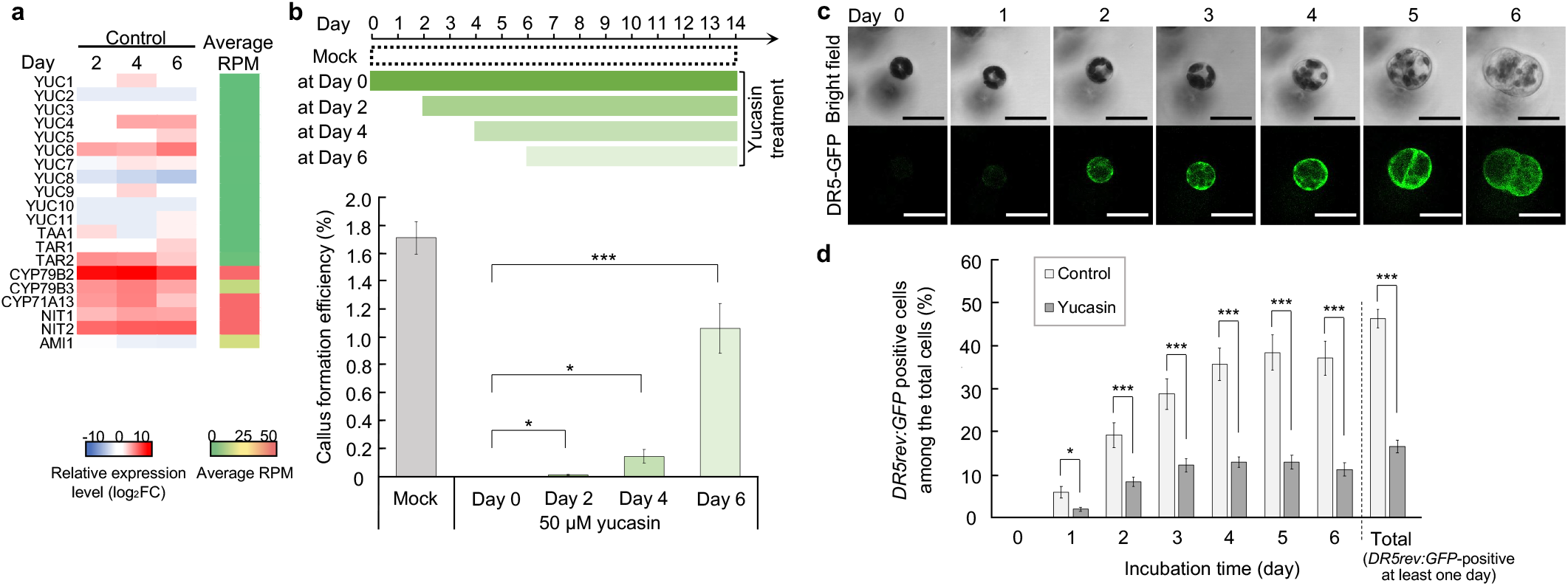
Early auxin biosynthesis is required to reinitiate cell division in leaf mesophyll protoplasts. **a**, Heat map of the transcriptional changes for representative IAA biosynthesis genes at Day 2, 4 and 6 in the control condition. Values (log_2_FC) are relative to Day 0. Average RPM indicates the overall expression level for each gene. **b**, Callus formation efficiency of WT protoplasts treated with 50 μM yucasin at different time points. Diagram shows the timing of yucasin treatment. Error bars represent standard error. n = 15 from 3 biological replicates. \**P* < 0.05, \*\*\**P* < 0.001 (two-tailed Welch’s *t*-test compared to yucasin treatment at Day 0). **c**, Time-lapse confocal microscopy images of a mesophyll protoplast that undergoes cell division at Day 5. Auxin response is visualized by *DR5rev:GFP* expression. **d**, Frequency of *DR5rev:GFP*-expressing (positive) protoplasts in the control or 50 μM yucasin condition among all tested *DR5rev:GFP* protoplasts. Error bars represent standard error. n = 9 for control and 7 for yucasin-treated protoplasts from 2 biological replicates. \*\*\**P* < 0.001 (two-tailed Student’s *t*-test or Welch’s *t*-test). Scale bars are 30 μm (**c**).

### Auxin biosynthesis is required to transcriptionally activate G2/M phase genes

To further reveal how auxin promotes resumption of the mitotic cell cycle, we compared expression patterns of core cell cycle regulators under control and yucasin conditions by RNA-seq (Fig. 4a and Supplementary Table 7). Previous studies have shown that freshly isolated leaf mesophyll protoplasts reside at the G1 phase and enter the S phase only upon phytohormone application^15^. Consistently, genes functioning during the G1 to S phase, including *D-type CYCLIN*s (*CYCD*s) and *MINICHROMOSOME MAINTENANCE 3* (*MCM3*), show upregulation from Day 2 in control condition. In agreement with our observation that the timing of initial cell division peaks around Day 6, genes functioning during the G2 to M phase, such as *B-type CYCLIN-DEPENDENT KINASE*s (*CDKB*s), *B-type CYCLIN*s (*CYCB*s), *CELL DIVISION CYCLE 20.1* (*CDC20.1*) and *CDC20.2*, are upregulated particularly at Day 4 and Day 6. Strikingly, in the yucasin condition, the expression of G1/S genes is comparable to that in the control condition, while many G2/M genes are clearly suppressed (Fig. 4a). This notion is further confirmed when a larger set of genes that show transcriptional activation in S phase or G2/M phase is examined^29^ (Supplementary Fig. 7a and Supplementary Table 7a-c), suggesting that auxin biosynthesis is required to transcriptionally activate the G2/M phase genes. We also observed similar transcriptional trends in the MB-3 condition (Fig. 4a, Supplementary Fig. 7a), supporting our hypothesis that one of the key downstream pathways regulated by histone acetylation is the IAA biosynthesis-dependent G2/M progression. Intriguingly, yucasin or MB-3 treatment has little impact on the transcription of photosynthetic genes (Supplementary Fig. 8 and Supplementary Table 2), suggesting that this aspect of cellular dedifferentiation is regulated by mechanisms independent from histone acetylation and IAA biosynthesis.

**Figure 4.**
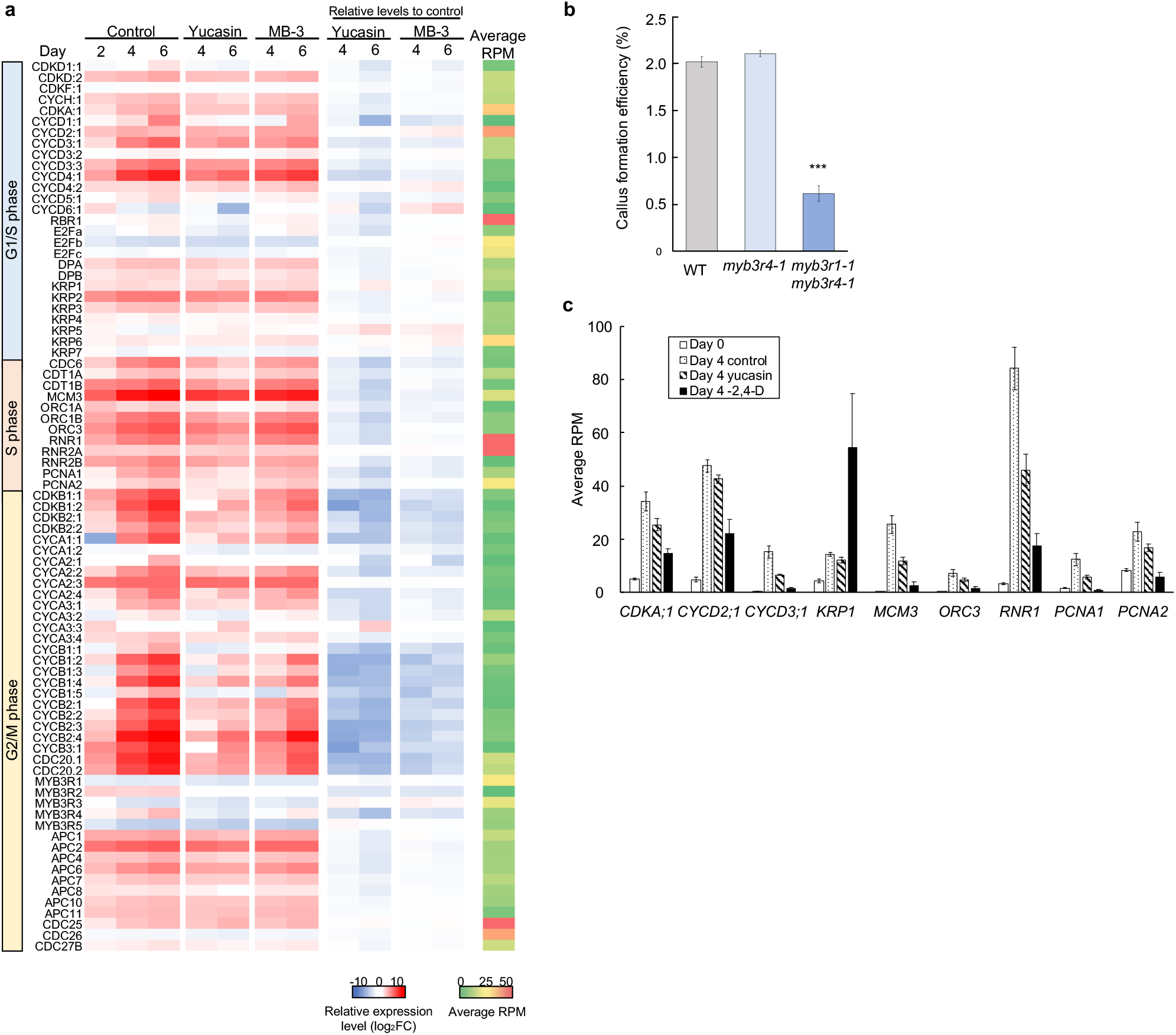
Auxin biosynthesis is required to transcriptionally activate the G2/M phase genes through MYB3R1 and MYB3R4. **a**, Heat map representing the transcriptional changes for genes encoding core cell cycle regulators at G1/S, S and G2/M phases at Day 2, 4 and 6. The left three columns show expression levels in the control, 50 μM yucasin and 50 μM MB-3 conditions as values normalized (log_2_FC) to Day 0. The ‘Relative levels to control’ columns show the normalized expression levels in the yucasin or MB-3 condition compared to the control condition for respective time points as relative values (log_2_FC). Average RPM indicates the overall expression level for each gene. Gene sets are selected based on Kalve *et al.*^47^. **b**, Callus formation efficiency of WT, *myb3r4-1* and *myb3r1-1 myb3r4-1* protoplasts. Error bars represent standard error. n = 20 from 4 biological replicates for the WT and *myb3r1-1 myb3r4-1* and n = 15 from 3 biological replicates for *myb3r4-1*. \*\*\**P* < 0.001 (two-tailed Student’s *t*-test compared to WT). **c**, Examples of the G1/S phase-related genes that are differently expressed in the - 2,4-D condition compared to the control and yucasin conditions at Day 4. Error bars represent standard error. n = 6 for Day 0 and the control condition at Day4 and n = 3 for the yucasin and - 2,4-D conditions at Day 4.

The identification of a number of G2/M genes transcriptionally targeted by auxin biosynthesis and histone acetylation suggests that some master regulators of these genes are involved in this regulation. Previous studies have shown that MYB3R4, together with MYB3R1, functions as a transcriptional activator of G2/M genes and their peak in activity at the G2/M phase is regulated at transcriptional and/or post-translational levels^29–32^. Indded, our RNA-seq data show that the expression of *MYB3R4* is sharply upregulated from Day 2 to Day 6 in control but not in either the yucasin or MB-3 condition (Fig. 4a), implying that auxin biosynthesis and histone acetylation regulate cell cycle progression by transcriptionally upregulating *MYB3R4*. To test whether these activator MYB3Rs regulate cell cycle reentry in protoplasts, we isolated protoplasts from *myb3r4-1* or *myb3r1-1 myb3r4-1* mutants and tested their callus formation efficiencies. As shown in Fig. 4b, *myb3r1-1 myb3r4-1* mutants, in which overall expression of G2/M genes are strongly repressed^30,31^, have severely impaired callus formation efficiency (Fig. 4b), indicating that these activator MYB3Rs are required for cell cycle reinitiation in protoplasts.

The discovery that auxin biosynthesis is critical for protoplast cell cycle reinitiation is surprising considering that PCIM contains exogenous auxin, 2,4-D, which is indispensable for initial cell division^33^ (data not shown). Previous studies have indicated that exogenous auxin regulates the G1/S transition at both transcriptional and post-translational levels in protoplasts^33,34^, raising the possibility that exogenous and endogenous auxin regulate different phases of the cell cycle to reinitiate cell division in protoplasts. To test this hypothesis, we compared the transcriptional activation of S phase and G2/M phase genes in control, yucasin and 2,4-D-omitted (- 2,4-D) culture conditions. Our data show that many of genes essential for the S phase progression, such as *CYCD2;1*, *CYCD3;1* and *MCM3*, are clearly suppressed in the - 2,4-D condition compared to the control and yucasin conditions, whereas G2/M phase genes show similar levels of hypo-activation in both the yucasin and - 2,4-D conditions (Fig. 4c, Supplementary Fig. 7b and Supplementary Table 7d-e). We also found that *KIP-RELATED PROTEIN 1* (*KRP1*), an inhibitor for CDKA/CYCD complexes, is strongly overexpressed in the - 2,4-D condition (Fig. 4c), further supporting transcriptional repression of the G1/S transition. These results therefore suggest that exogenous and endogenous auxin have distinct roles during the initial division of protoplasts, where they transcriptionally activate the S phase and G2/M phase progression, respectively.

### ARF7/ARF19 and IAA3/IAA18-mediated auxin signalling pathway regulates cell cycle reinitiation in protoplasts

Having established the central roles of auxin in protoplast reprogramming, we next sought to investigate which auxin signalling pathways participate in this regulation. As reported previously, auxin is also critical for other forms of cellular reprogramming in plants, including pluripotent callus formation from explants in tissue culture^4^. In Arabidopsis auxin promotes cell cycle reactivation in pericycle cells on callus induction medium (CIM), and this is mediated by ARF7, ARF19, IAA14 and LATERAL ORGAN BOUNDARIES DOMAINs (LBDs)^35–37^. Our RNA-seq data show that many of these auxin signalling regulators are sharply upregulated before initial cell division of protoplasts (Fig. 5a and Supplementary Table 8), implying that they also participate in cell cycle reinitiation from differentiated leaf cells. A loss-of-function mutant for ARF7, *non-phototrophic hypocotyl 4-1* (*nph4-1*)^38^, indeed displays defects in the auxin response in mesophyll protoplasts^39^ and consistently, single or double mutants for *ARF7* and *ARF19* are impaired in callus formation from protoplasts (Fig. 5b, Supplementary Fig. 9a). Intriguingly, however, a gain-of-function mutant for *IAA14*, *solitary root-1* (*slr-1*)^40^, and two loss-of-function mutants for *LBD*s, *pLBD16:LBD16-SRDX* and *lbd16-1 lbd18-1 lbd33-1*^41^ make callus from protoplasts with the similar efficiency as the WT, despite displaying severe defects in callus formation from hypocotyl explants in tissue culture^35^ (Fig. 5b-c, Supplementary Fig. 10). These results indicate that reprogramming from differentiated leaf cells is regulated by distinct auxin signalling pathways. To further investigate the auxin signalling components responsible for protoplast reprogramming, we selected other *Aux/IAA* candidates that have expression levels comparable with those of ARF7 and/or ARF19 in leaves and show strong binding to these ARFs^42^ (Supplementary Fig. 9b and Supplementary Table 9). Among them, we found that gain-of-function mutants for IAA3 and IAA18, *suppressor of hy2-101* (*shy2-101*)^43^ and *crane-2*^44^, respectively, strongly inhibit callus formation from protoplasts while a gain-of-function mutation in *IAA7*, *auxin resistant 2-1* (*axr2-1*)^45^ does not cause obvious defects (Fig. 5d). Collectively, our results suggest that IAA3 and IAA18, together with ARF7 and ARF19, mediate the auxin response in protoplasts, which drives cell cycle reinitiation.

**Figure 5.**
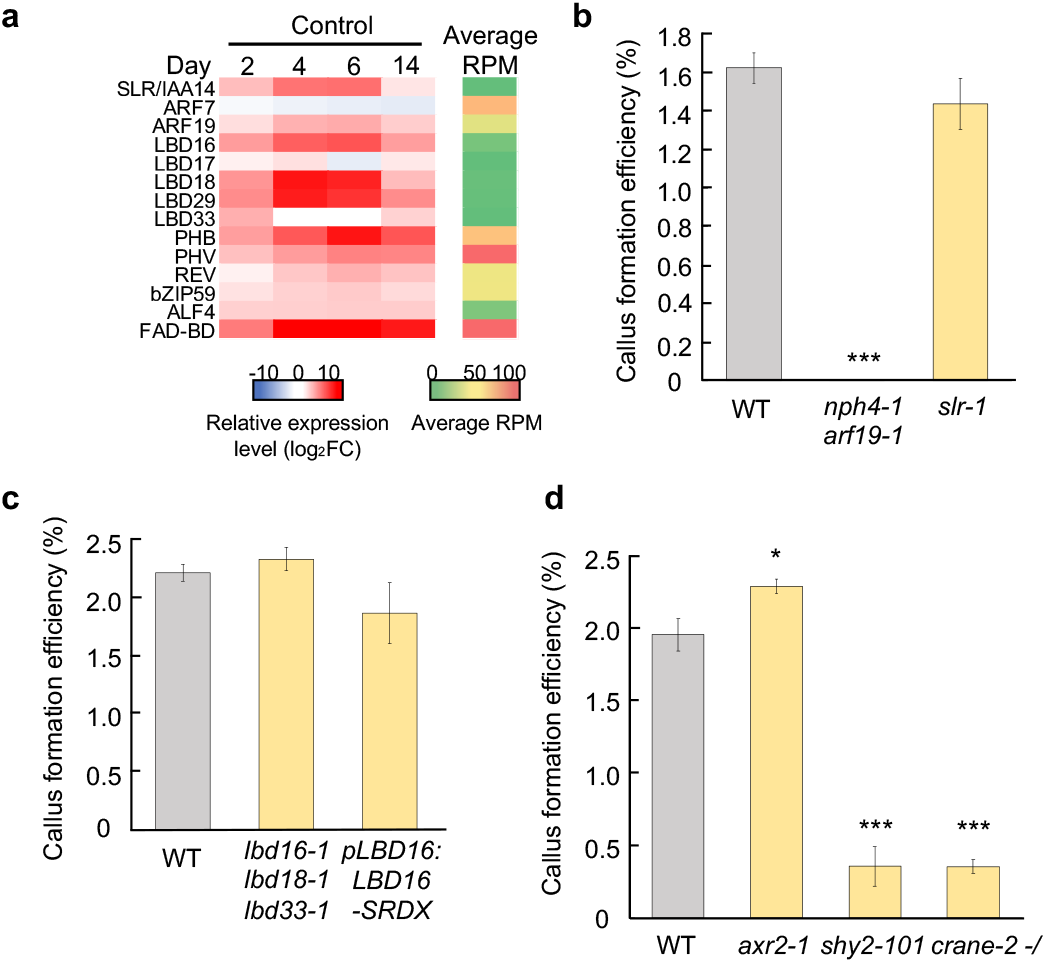
ARF7/ARF19 and IAA3/IAA18-mediated auxin signalling pathway drives cell cycle reinitiation in leaf mesophyll protoplasts. **a**, Heat map representing the transcriptional changes for genes implicated in auxin-induced callus formation in tissue culture. Expression levels at Day 2, 4 and 6 in the control condition are shown as values normalized (log_2_FC) to Day 0. Average RPM indicates the overall expression level for each gene. Gene sets are taken from Ikeuchi *et al.*^6^. **b**, Callus formation efficiency of WT, *nph4-1 arf19-1* and *slr-1* protoplasts. Error bars represent standard error. n = 25 from 5 biological replicates for the WT and 20 from 4 biological replicates for the others. \*\*\**P* < 0.001 (two-tailed Student’s *t*-test or Welch’s *t*-test compared to WT). **c**, Callus formation efficiency of WT, *lbd16-1 lbd18-1 lbd33-1* and *pLBD16:LBD16-SRDX* protoplasts. Error bars represent standard error. n = 20 from 4 biological replicates for the WT and 15 from 3 biological replicates for the others. No statistical difference was detected (two-tailed Student’s *t*-test or Welch’s *t*-test compared to WT). **d**, Callus formation efficiency of WT, *shy2-101*, *axr2-1* and *crane-2* protoplasts. Since heterozygous and homozygous *crane-2* are morphologically indistinguishable, both genotypes were mixed for protoplast preparation. Error bars represent standard error. n = 25 from 5 biological replicates for the WT and 15 from 3 biological replicates for the others. \**P* < 0.05, \*\*\**P* < 0.001 (two-tailed Student’s *t*-test or Welch’s *t*-test compared to WT).

## Discussion

In this study we demonstrate that one of the key factors that permit reprogramming of differentiated plant cells is the activation of auxin biosynthesis. We show that endogenously produced IAA is required to increase auxin response in protoplasts, thereby inducing the expression of G2/M genes to complete cell division (Fig. 6). Notably, auxin biosynthesis is critical specifically for their first cell division while it is not essential for successive divisions (Fig. 3b). Since differentiated cells should be equipped with mechanisms to prevent ectopic cell proliferation^46^, it is plausible that cell cycle reinitiation in protoplasts requires some unique mechanisms in addition to those functioning during successive cell proliferation, with the latter more similar to those at play in normal development. Previous studies, for instance, have shown that differentiating leaf cells accumulate regulators, such as DA1 and MEDIATOR 25 (MED25), that repress proliferation and promote cellular growth^47^. In parallel, they shut down the expression of many cell cycle activators such as *CYCD4*s and *ANAPHASE COMPLEX 10* (*APC10*) through epigenetic mechanisms^46,48^. Differentiating cells, additionally, develop physical properties such as thickened cell walls and enlarged vacuoles that may inhibit cell division. Restarting the cell cycle should therefore require the removal of these negative factors and/or induction of some potent activators that can drive cell cycle reentry. In the case of mesophyll protoplasts, auxin biosynthesis induces the transcription of G2/M phase genes likely through MYB3R4 and MYB3R1 (Fig. 4), suggesting that these MYB3Rs serve as key re-activators of cell division. These MYB3Rs appear to be dispensable for mesophyll cell proliferation in intact leaves^31^, so they might be required to induce cell division specifically in differentiated cells where other factors promoting cell cycle progression are possibly inactivated. How auxin biosynthesis regulates MYB3R4/1-mediated pathways is an important question, and its mechanistic details should be further investigated in future studies. Our data suggest that auxin biosynthesis transcriptionally activates MYB3R4 (Fig.4a), but it, as well as MYB3R1, might be also subject to post-translational regulation since phosphorylation and temporal nuclear shuttling of MYB3Rs are central for their functions during mitosis^32,49^.

**Figure 6.**
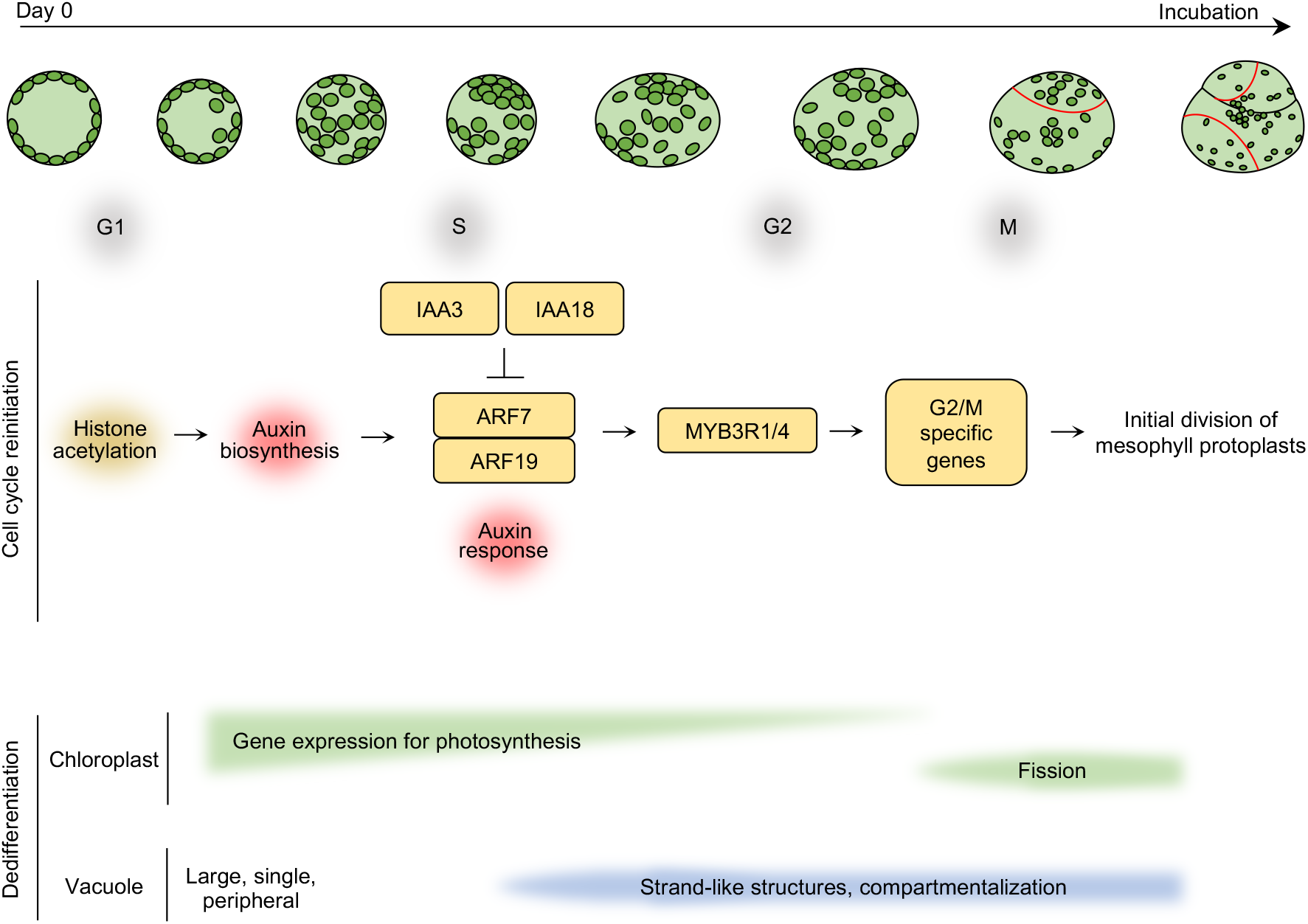
Hypothetical model describing the molecular mechanism of developmental reprogramming in leaf mesophyll protoplasts. Freshly isolated Arabidopsis leaf mesophyll protoplasts undergo dramatic developmental reprogramming and produce pluripotent callus when cultured in the presence of auxin and cytokinin. Histone acetylation is required to reinitiate the mitotic cell cycle and one of the key downstream pathways regulated by histone acetylation is the transcription of auxin biosynthesis genes. Endogenously produced auxin, IAA, in turn activates auxin response in protoplasts, possibly in an ARF7/ARF19 and IAA3/IAA18-dependent manner. Endogenous auxin is responsible for the transcriptional upregulation of G2/M genes to complete the initial cell division, and such cell cycle regulation likely involves auxin-dependent induction of MYB3R4. MYB3R4 and MYB3R1 might be also regulated at the post-translational level. Another key feature of developmental reprogramming that protoplasts undergo at an early stage of culture is the cellular dedifferentiation, which is marked, for instance, by drastic downregulation of photosynthetic genes and upregulation of chloroplast fission genes. These transcriptional changes are independent of histone acetylation and auxin biosynthesis. In addition, a large, single vacuole, typical of differentiated cells, becomes compartmentalized and starts to resemble what is found in proliferating or elongating cells.

We predict that exogenously supplied 2,4-D serves as another strong driver of cell cycle reinitiation in protoplasts and interestingly, our data suggest that 2,4-D and endogenously produced IAA activate distinct phases of the cell cycle to reinitiate protoplast division (Supplementary Fig. 7b). This is consistent with earlier reports that exogenous 2,4-D is required during the initial steps of protoplast culture^33^, while IAA biosynthesis is activated later on (Fig. 2h; Pasternak et al., 2002). To further clarify how these two auxins cooperate to promote cell cycle reinitiation, it will be helpful to investigate whether the ARF7/19- and IAA3/18-mediated pathway transduces 2,4-D- and/or IAA-induced signalling. Alternatively, it is possible that stimulation of the IAA biosynthetic pathway itself plays a crucial role in protoplast division. For instance, the YUC cofactor flavin adenine dinucleotide (FAD) is oxidized and produces hydrogen peroxide (H_2_O_2_) when the IAA precursor indole 3-pyruvate (IPA) is deficient^50^. H_2_O_2_ has been demonstrated to either promote or inhibit cell cycle reinitiation in protoplasts depending on the timing of its production and subcellular localization^12,13^. Since *YUC* genes also need to be upregulated at an appropriate dose and time during protoplast culture (Fig. 2h), activation of the IAA biosynthetic pathway may contribute to protoplast reprogramming through the production of its byproducts such as H_2_O_2_. It is also worth considering the possibility that overall regulation of indole-containing compound metabolism affects protoplast reprogramming, since our RNA-seq data show the striking upregulation of the indole metabolism-related genes that may not act in the IAA biosynthetic pathway (Fig. 2e; Supplementary Table 4). Investigating the potential contribution of these non-hormonal metabolites may help uncover a new regulatory mechanism underlying plant cellular reprogramming.

## Methods

### Plant materials and growth conditions

Arabidopsis thaliana (L) Heynh. ecotype Columbia-0 (Col-0) was used as the wild type. For phenotypic analyses, *hag1-1* (SALK_150784), *hag1-2* (SALK_030913), *hag2* (SALK_051832), *hag3-1* (GABI_555H06), *hag3-2* (SALKseq_060819), *ham1* (SALK_027726), *ham2* (SALK_106046), *hac1* (SALK_080380), *hac2* (SALK_049434), *hac4* (SALK_045791), *hac5* (SAIL_49_D10), *hac12* (SALK_012469), *haf1-1* (SAIL_256_D10), *haf1-2* (SALK_110848), *haf2* (SALK_110029), *yucQ* (GABI_376G12 x CSHL_GT6160 x SALK_059832 x SM_3_23299 x SALK_762_D07)^25^, *myb3r4-1* (SALK_059819), *myb3r1-1 myb3r4-1* (SALK_018482 x SALK_059189)^30^, *arf7-1* (SALK_040394), *arf19-4* (SALK_009879), *nph4-1 arf19-1*^38^, *arf7-2 arf19-5*^41^, *slr-1*^40^, *lbd16-1 lbd18-1 lbd33-1* (SALK_095791 x SALK_038125 x SAIL_95_H10)^41^, *axr2-1*^45^, *shy2-101*^43^ and *crane-2*^44^ were used. *cyp79B2 cyp79B3* mutants were generated by crossing *cyp79B2* (SALK_130570C) with *cyp79B3* (SALKseq_066556). The *pVHP1:VHP1-mGFP*^21^ and *DR5rev:GFP* lines^28^ were previously described. All these mutants and transgenic plants were in Col-0 background. Seeds were sterilized with 70 % ethanol for 1 min and 20 % chlorine bleach (Kao) for 10 min, and then rinsed by autoclaved water for 3 times. After being soaked in water at 4□ for 2 to 4 days, seeds were sown on gemination medium (GM) (Supplementary Table 1) with the density of 33 seeds per 90-mm diameter and 25-mm thick polystyrene dish (Kord-Valmark). Plants were grown at 22□, under continuous light (30 to 40 μmol /m^2^ /s) in a growth chamber (Sanyo).

### Transgenic plants

To construct the *pCAB3:H2A-eGFP* vector, *CAB3* promoter, 1,537 bp upstream of the ATG start codon, was amplified from genomic DNA by PCR. The PCR products were purified and cloned into the *pDONRP4P1r* vector (Thermo Fisher Scientific). The promoter fragment was then assembled with GAL4 into the *pB-9FH2A-UAS-7m24GW* destination vector in a multi-site gateway reaction to create an activator line construct using the LR Clonase II+ (Thermo Fisher Scientific). This destination vector contains a HISTONE 2A-6 (H2A) encoding sequence fused to eGFP and driven by the repetitive UAS promoter, as described by Fendrych et al. (2014)^51^. To construct the *pHAG1:HAG1-GFP* vector, *HAG1* promoter, 1,987 bp upstream of the ATG start codon, was amplified from genomic DNA, purified and cloned into the *pDONRP4P1r* vector (Thermo Fisher Scientific). *HAG1* coding sequence was amplified from complementary DNA (cDNA) by PCR, purified and cloned into the *pENTR/D-TOPO* vector (Thermo Fisher Scientific). The promoter and coding sequence fragments were then assembled with GFP into the *R4pGWB504* destination vector in a multi-site gateway reaction using the Gateway LR Clonase □ (Thermo Fisher Scientific). To construct the *XVE-YUC1* vector, *YUC1* coding sequence was amplified from cDNA by PCR. The PCR products were purified and cloned into *pENTR/D-TOPO* vector (Thermo Fisher Scientific). The cDNA fragments were then cloned into the modified *pER8* plasmid containing a Gateway cassette (*pER8-GW*)^52^ using the Gateway LR Clonase □ (Thermo Fisher Scientific). For plant transformation, the plasmids were introduced into *Agrobacterium tumefaciens* (strain GV3101) by electroporation and transformed into *Arabidopsis* Col-0 WT plants (for *pCAB3:H2A-eGFP* and *XVE-YUC1*) or *hag1-1* heterozygous plants (for *pHAG1:HAG1-GFP*) by the floral dip method^53^. As for *pHAG1:HAG1-GFP/hag1-1* plants, after the selection of single insertion lines of *pHAG1:HAG1-GFP* at T2, the lines homozygous for *hag1-1* mutation were further selected by PCR genotyping. Lines homozygous both for *pHAG1:HAG1-GFP* and *hag1-1* were then selected at T3. A list of primers used for PCR is described in Supplementary Table 10.

### Chemical compounds

2,4-dichlorophenoxyacetic acid (2,4-D) (Cas 94-75-7, Sigma), C646 (Cas 328968-36-1, Sigma)^54^, β-estradiol (ED) (Cas 50-28-2, Wako), garcinol (Cas 78824-30-3, Focus biomolecules), kinetin (Kin) (Cas 525-79-1, SIGMA), L-kynurenine (Kyn) (Cas 2922-83-0, TCI)^56^, γ-butyrolactone (MB-3) (CAS 778649-18-6, Abcam)^57^, thidiazuron (TDZ) (Cas 51707-55-2, Wako), 5-(4-chlorophenyl)-4H-1,2,4-triazole-3-thiol (yucasin) (CAS 26028-65-9, Wako)^58^, 6-(γ,γ-Dimethylallylamino)purine (2-iP) (CAS 2365-40-4, Sigma), were dissolved in dimethyl sulfoxide (DMSO) and sterilized by filtration. Indole-3-acetic acid (IAA) (Cas 6505-45-9, Wako) was dissolved in ethanol and sterilized by filtration. Fluorescein diacetate (FDA) (Cas 596-09-8, DOJINDO) was dissolved in DMSO and used without sterilization. All chemicals were stored at -20□.

### Protoplast isolation and callus induction

Protoplasts were isolated following Damm and Willmitzer^59^ with modifications. All processes were performed at room temperature (22 to 25□) unless otherwise specified. 1st to 5th rosette leaves of 23 or 24 days after sowing (DAS) plants were aseptically harvested, and after carefully removing petioles, chopped into strips with a scalpel (Akiyama) in 0.5 M mannitol. After 1-hour maceration in 0.5 M mannitol under dim light, leaf strips from 100 to 150 leaves were transferred into a 35-mm polystyrene petri dish (FALCON) and soaked in 3 mL of Digestion Cocktail (Supplementary Table 1). For cell wall digestion, leaf strips were gently shaken horizontally on Shake-LR (TAITEC) for 3 hours in the dark. Protoplasts were then separated from leaf tissues by filtration through a 40 μm nylon Cell Strainer (FALCON) with gentle pipetting using a Komagome type pipette (IWAKI). The filtrate was collected into a 12 mL culture tube with a conical bottom (Simport), diluted with 1/2 volume of 0.2 M CaCl_2_ and centrifuged for 5 min at 60 g in a swinging bucket rotor (CF16RN, HITACHI), with the slowest acceleration and no brake. The pellet was resuspended in 5 mL of Wash Medium 1 (Supplementary Table 1) and centrifuged for 3 min at 40 g. The resulting pellet was resuspended in 5 mL of Wash Medium 2 (Supplementary Table 1) and centrifuged again. The pellet was subsequently resuspended in 5 mL of 0.5 M mannitol and centrifuged again. The protoplasts were finally resuspended in fresh 0.5 M mannitol and placed on ice for 40 min to 1 hour under dim light. To test the viability of fresh protoplasts, FDA was added to an aliquot of protoplast solution at the final concentration of 1 mg/L and incubated for 10 min. The stained cells were mounted onto a microscope slide and observed with a fluorescence microscope (BX51, OLYMPUS). After calculating cell density with a hemocytometer (Sunlead Glass Corp.), the protoplasts were warmed to room temperature and their density was adjusted to 4.8 to 5.0 ×10^5^ cells /mL with 0.5 M mannitol.

Isolated protoplasts were cultured using the protocol modified from Damm and Willmitzer^59^, Masson and Paszkowski^60^ and Chupeau *et al.*^61^. For embedding protoplasts in the sodium alginate gel, the protoplast solution was gently mixed with an equal volume of Sodium Alginate Solution (Supplementary Table 1) and 200 μL of this solution was then dropped onto CaCl_2_ Plate (Supplementary Table 1) using a truncated tip. Sodium alginate gels, each containing approximately 4.8 to 5.0 ×10^4^ cells, were solidified into a piece of gel for 1 hour at room temperature under dim light. Five pieces of gels were subsequently transferred into a well of 6-well microplates (IWAKI) or a 35-mm petri dish (FALCON) filled with 4 mL of protoplast callus induction medium (PCIM) (Supplementary Table 1). The plates and dishes were then sealed with surgical tape and parafilm, respectively, and embedded protoplasts were aseptically cultured at 22□ in the dark.

Callus formation efficiency was evaluated at Day 14 unless otherwise specified. Calli embedded in each sodium alginate gel were counted under a dissection microscope (M165 FC, Leica) at 2.5× magnification. Callus formation efficiency was calculated as a percentage of the number of calli in the number of protoplasts initially contained in one sodium alginate gel. The reproducibility was confirmed in at least 3 biological replicates. As the control experiments in chemical treatment assays, protoplasts of corresponding genotypes were treated with the equivalent volumes of solvents (DMSO or ethanol) to confirm that their callus formation efficiencies are not affected by these treatments (data not shown in some figures).

### Induction of shoot formation from protoplast-derived callus

After incubating in PCIM for 14 days, calli-containing gels were transferred to 4 mL of callus growing medium (CGM) (Supplementary Table 1) and cultured at 22□ under continuous dim light (10 to 12 μmol /m^2^ /s). CGM was refreshed every 2 weeks. For induction of *de novo* shoot formation, calli were isolated from the gels by shaking them in Citrate Solution (Supplementary Table 1) for 1 to 2 hours, which chelates calcium ions and dissolves the gel. Isolated calli were washed twice by liquid Gamborg B5 medium and cultured on shoot induction medium (SIM) (Supplementary Table 1) at 22□ under continuous light (22-28 µmol /m^2^ /s).

### Callus induction from etiolated hypocotyls

To induce callus from tissues, seeds were sown on GM for Tissue Culture (Supplementary Table 1) and grown at 22□ in the dark for 7 days. Hypocotyls of etiolated seedlings were excised into around 5-mm-long explants using a scalpel and incubated on Callus Induction Medium (CIM) (Supplementary Table 1) for 21 days at 22□ under continuous light (22-28 μmol /m^2^ /s). The reproducibility was confirmed in 2 biological replicates.

### Live imaging of protoplast reprogramming

The microscopic observation method for tracking identical protoplasts was developed based on Hall *et al.*^62^ and Dovzhenko *et al.*^63^. To immobilize the protoplast-containing gels on the bottom of a 35-mm petri dish, a polypropylene grid (1.7 ×1.7 mm mesh size, NIP), cut into 11 ×11 meshes, were placed a CaCl_2_ Plate and the protoplast solution was solidified on the mesh. The gel was subsequently fixed to the bottom of a petri dish using the attached grid and additional 4 gels were placed to the same dish to keep the number of protoplasts within a well constant as in our standard culture condition. These gels were cultured with 4 mL of PCIM at 22□ in the dark.

Time-lapse imaging was performed using a confocal microscope (TCS-SP5, Leica) with a water immersion lens (HC FLUOTAR L 25×/ 0.95 W VISIR, Leica). The protoplast-containingpetri dishes were placed on the microscopic stage at each time of observation. Region of Interests were manually tracked based on the coordinates (x, y, z) recorded on the first day. Z-stacked images were taken at 6 μm intervals to maximize the number of protoplasts that can be tracked. Each observation was performed within 1 hour per dish to avoid excessive light exposure of protoplasts. Images were processed and analyzed with Fiji ver. 2.0.0 (https://imagej.net/Fiji) and Microsoft Excel. The reproducibility was confirmed in at least 2 biological replicates.

### Transcriptome analysis

RNA was extracted from ∼1 g (in fresh weight) of rosette leaves or 1.0 to 1.5 ×10^6^ protoplasts that were prepared from 23 DAS plants. Freshly isolated protoplasts were collected as pellet in a 2-mL tube by centrifugation for 1 min at 3,200 g (MX-305, TOMY) and stored at -80□. To collect cultured protoplasts embedded in the sodium alginate gel, 25 gels per sample were soaked into 25 mL of Citrate Solution and protoplasts were released by gentle shaking for 1 hour. Three biological replicates were prepared for each time point. Total RNA was isolated using the RNeasy plant mini kit (Qiagen). Isolated RNA was then subjected to library preparation using the Kapa stranded mRNA sequencing kit (Kapa Biosystems) with NEBNext Multiplex Oligos for Illumina (New England Biolabs) as adapters and Agencourt AMPure XP (Beckman Coulter) beads instead of KAPA Pure Beads. Single-end sequencing was performed on an Illumina NextSeq500 platform. Mapping was carried out using Bowtie ver. 0.12.9. Over 50 % of the reads were uniquely mapped to the TAIR10 Arabidopsis genome, resulting in 5 to 17 million mapped reads per sample. Differentially expressed genes were identified using the edgeR package^64^ on R/Bioconductor (https://www.r-project.org/) after normalization of total read counts with Trimmed Mean of M-values (TMM) method. Differentially expressed genes were defined as those that showed │log_2_FC│> 2 in transcript levels (*p*-value < 0.01 and FDR < 0.01). Gene ontology (GO) analyses were performed by PANTHER GO Enrichment Analysis^65–67^ (http://geneontology.org) with PANTHER Overrepresentation Test (Released 20210224). Genes were annotated following GO Ontology database (Released 2021-02-01) and categorized by Biological Processes. Venn diagrams were drawn with BioVenn^68^ (http://www.biovenn.nl) and processed with Inkscape ver. 0.92.4 (https://inkscape.org). Heat maps were drawn using Color Scale tool in Microsoft Excel.

### Reverse transcription-quantitative polymerase chain reaction (RT-qPCR)

500 ng of the extracted total RNA were subjected to the first-strand cDNA synthesis with the primescript RT reagent kit (Takara). Quantitative real-time PCR was performed with Thunderbird SYBR qPCR mix (Toyobo) in Stratagene MX3000P real-time qPCR system (Agilent Technologies). The transcript levels were calculated with standard curve method and normalized by that of internal control *PROTEIN PHOSPHATASE 2A-3* (*PP2A3*)^69^. A set of primers used for qPCR is described in Supplementary Table 10.

## Acknowledgements

We thank Munetaka Sugiyama (the University of Tokyo) for technical advice on protoplast culture, Masaki Ito (Kanazawa University) for the seeds of MYB mutants and Hidehiro Fukaki (Kobe University) for the seeds of ARF and Aux/IAA mutants. This work was supported by a grant from the Ministry of Education, Culture, Sports, and Technology of Japan to K.S. (20H03284 and 20H05911) and Grant-in-Aid for JSPS Fellows to Y.S. (20J20380). Y.S. is supported by JSPS DC research fellowship.

## Author contributions

Y.S. and K.S. conceived the project. Y.S. and K.S. designed the experiments, and Y.S. conducted most of genetic and cell biological analyses except for RNA sequencing and mapping which were performed by T.S. A.K. generated *pHAG1:HAG1-GFP/hag1-1* plants. S.P. and L.D.V. generated *pCAB3:H2A-eGFP* plants. S.S. and M.M. provided *pVHP1:VHP1-mGFP* seeds. Y.S. and K.S. wrote the manuscript with help from the co-authors.

## Competing interests

The authors declare no competing interests.

## Additional information

Supplementary data are available for this paper online. Correspondence and requests for materials should be addressed to K.S.

## Supplementary Figure legends

**Supplementary Figure 1** │ **Cell cycle reinitiation and callus formation described with a newly established culture system and time-lapse confocal microscopy. a**, Bright-field and fluorescence microscopy images of freshly isolated WT protoplasts. FDA staining indicates that 98.66 ± 0.19 % (standard error, n = 10) of freshly isolated protoplasts are viable. **b**, Callus formation efficiency of WT protoplasts. Average efficiency is 1.98 ± 0.33%. n = 240 from 48 biological replicates. Each dot shows callus formation efficiency calculated from one gel. In the box plot, the median is represented by a black line and the upper and lower quartiles are represented by the upper and lower ends of the box respectively. **c**, Bright-field and fluorescence microscopy images of protoplasts freshly isolated from plants carrying a mesophyll cell-specific marker *pCAB3:H2A-eGFP*. Quantitative analysis indicates that 94.63 ± 0.51% of freshly isolated protoplasts show distinct nuclear-localized H2A-eGFP expression (standard error, n = 15 from 2 biological replicates). **d**, Diagram showing the time-lapse confocal microscopy procedure used to track individual protoplasts. **e,** Classification of 164 protoplasts carrying *pCAB3:H2A-eGFP* that underwent cell division by Day 10 based on H2A-eGFP expression and appearance at Day 0. **f**, Two examples of *pCAB3:H2A-eGFP* protoplasts that are H2A-eGFP-positive and mesophyll cell-like at Day 0. Both protoplasts divided by Day 10. **g,** A *pCAB3:H2A-eGFP* protoplast that is H2A-eGFP-negative and mesophyll cell-like at Day 0. The protoplast divided by Day 10. **h,** A *pCAB3:H2A-eGFP* protoplast that is H2A-eGFP-negative and guard cell-like at Day 0. The protoplasts divided by Day 10. **i,** Heat maps representing cell size dynamics of protoplasts that reinitiate cell division between Day 4 and Day 9. Each row shows cell diameter (left panel) and change in cell diameter comparted to Day 0 (right panel) for individual protoplasts isolated from WT, *DR5rev:GFP* or *pVHP1:VHP1-mGFP* plants. Among 94 protoplasts tested in this experiment, 80 cells underwent cell elongation before cell division. **j,** Timing of cell elongation and cell division for 80 cells from **i**. **k,** Duration of cell elongation (right panel) for 80 cells from **i**. Scale bars are 100 μm (**a**), 100 μm (**d**) and 20 μm (**f** to **h**).

**Supplementary Figure 2** │ **Time-lapse confocal microscopy images of leaf mesophyll protoplasts that did or did not undergo cell division. a**, Pie chart showing the percentages of protoplasts that underwent cell division (divided), those that elongated without cell division (elongated), those that expanded without cell division (expanded) and those that shrunk or displayed no shape changes (others) among 902 protoplasts isolated form *DR5rev:GFP* plants and used for time-lapse confocal microscopy. **b**, Representative time-lapse images of elongated, expanded and other protoplasts from Day 0 to Day 7. **c,** Another set of time-lapse images of a protoplast that underwent cell division. **d,** Time-lapse images of a protoplast that elongated without cell division. **e**, Time-lapse images of a protoplast that expanded without cell division. In **c** to **e**, vacuolar morphology is visualized by VHP1-mGFP. The double-headed arrow indicates the direction of cell elongation and arrowheads mark the plane of initial cell division. The white arrow highlights the initial appearance of vacuolar strand-like structures. Scale bars are 30 μm.

**Supplementary Figure 3** │ **Transcriptional changes of chloroplast-related genes during protoplast reprogramming.** Heat map representing the transcriptional changes for genes implicated in chloroplast biogenesis and function. The leftmost column shows relative expression levels (log_2_FC) in protoplasts at Day 0 compared to 23 DAS leaves. The second column shows relative expression levels (log_2_FC) in cultured protoplasts at Day 2, 4, 6 and 14 compared to Day 0. Average reads per million (RPM) indicates the overall expression level for each gene. The right column shows genes significantly upregulated (orange) or downregulated (blue) during leaf development based on Andriankaja et al. (2012). Gene sets were selected based on annotations at The Arabidopsis Information Resource (TAIR) (https://www.arabidopsis.org/index.jsp).

**Supplementary Figure 4** │ **Roles of histone acetyltransferases in protoplast cell cycle reinitiation. a**, Callus formation efficiency of WT protoplasts treated with 50 μM MB-3 at different time points. Diagram shows the timing of MB-3 treatment. Error bars represent standard error. n = 15 from 3 biological replicates. n.s. not significant (two-tailed Student’s *t*-test or Welch’s *t*-test compared to MB-3 treatment at Day 0). **b**, Callus formation efficiency of WT protoplasts treated with C646 or garcinol. Error bars represent standard error. n = 30 from 6 biological replicates for the WT control and n = 15 from 3 biological replicates for all others. \*\*\**P* < 0.001 (two-tailed Student’s *t*-test or Welch’s *t*-test compared to WT control). n.d. not determined. **c**, Callus formation efficiency of protoplasts isolated from WT and HAT mutants. Error bars represent standard error. n = 45 from 9 biological replicates for the WT and n = 10 or 15 from 2 or 3 biological replicates for mutants. \**P* < 0.05, *** *P* < 0.001 (two-tailed Student’s *t*-test or Welch’s *t*-test compared to WT). **d**, Callus formation efficiency of WT, *hag1-2*, *hag3-2* and *haf1-2* protoplasts. Error bars represent standard error. n = 25 from 5 biological replicates for the WT and n = 15 from 3 biological replicates for all others. *** *P* < 0.001 (two-tailed Welch’s *t*-test compared to WT). **e**, Callus formation efficiency of WT and *pHAG1:HAG1-GFP*/*hag1-1* protoplasts. Error bars represent standard error. n = 15 from 3 biological replicates. No statistical difference was detected (two-tailed Student’s *t*-test compared to WT).

**Supplementary Figure 5 │ Roles of auxin biosynthesis in protoplast cell cycle reinitiation. a,** Callus formation efficiency of Kyn-treated WT protoplasts. Error bars represent standard error. n = 30 from 6 biological replicates for the WT control and n = 15 to 25 from 3 to 5 biological replicates. \*\*\**P* < 0.001 (two-tailed Student’s *t*-test or Welch’s *t*-test compared to WT control). **b**, Callus formation efficiency of WT protoplasts incubated with PCIM supplemented with 0 to 1,000 μg/L IAA. Error bars represent standard error. n = 10 to15 from 2 to 3 biological replicates. \*\**P* < 0.01, \*\*\**P* < 0.001 (two-tailed Student’s *t*-test or Welch’s *t*-test compared to WT control). **c**, β-estradiol-inducible expression of *YUC1* in *XVE-YUC1* plants. WT and *XVE-YUC1* seeds were sown in liquid half-strength MS medium and incubated at 22□ under light with rotation on a Shake-LR (TAITEC) for 10 days. The plants were then treated with different concentrations of ED or the equivalent volumes of DMSO and grown for another 24 hours. The expression levels of *YUC1* are normalized by those of the internal control *PP2A3* and shown as relative values compared to WT given the same treatments. The *YUC1* expression in WT remained constant throughout the treatments. The numbers above each bar are average expression levels. Error bars represent standard error (n = 6).

**Supplementary Figure 6** │ **Correlation between auxin response and the morphological changes during reprogramming of protoplasts. a**, Frequency of the *DR5rev:GFP*-expressing (positive) protoplasts among 902 total cells, 45 divided cells, 221 elongated cells, 89 expanded cells and 547 other cells. *DR5rev:GFP* protoplasts were incubated in the control condition. \**P* < 0.05 (two-tailed Student’s *t*-test or Welch’s *t*-test compared to ‘Total’ at the same time point). **b**, Heat maps representing cell size dynamics of 417 *DR5rev:GFP*-positive and 485 *DR5rev:GFP*-negative protoplasts in the control condition. **c,** Heat maps representing cell size dynamics of 124 *DR5rev:GFP*-positive and 606 *DR5rev:GFP*-negative protoplasts in the 50 μM yucasin condition. In **b** and **c**, each row shows cell diameter (left panel) and change in cell diameter comparted to Day 0 (right panel) for individual *DR5rev:GFP* protoplasts.

**Supplementary Figure 7** │ **Roles of histone acetylation, auxin biosynthesis and exogenous 2,4-D in regulating expression of cell cycle genes during protoplast reprogramming. a**, Heatmap representing the transcriptional changes for 172 genes specifically expressed at S phase and 185 genes specifically expressed at G2/M phase. The left three columns show expression levels in the control, 50 μM yucasin, and 50 μM MB-3 condition as values normalized (log_2_FC) to Day 0. The ‘Relative levels to control’ columns show the normalized expression levels in the yucasin or MB-3 condition compared to the control condition for respective time points as relative values (log_2_FC). Average RPM indicates the overall expression level for each gene. Gene sets are taken from Kobayashi *et al.*^29^. Two genes in the original gene set were omitted from the heatmap of S phase genes since they were not expressed in all samples (see **Supplementary Table 7b**). **b,** Heatmap representing the transcriptional changes for the same gene sets as in **a**. The left three columns show expression levels at Day 4 in the control, 50 μM yucasin, and 2,4-D-omitted (named ’- 2,4-D’) condition as values normalized (log_2_FC) to Day 0. The ‘Relative levels to control’ columns show the normalized expression levels in yucasin or - 2,4-D condition compared to the control condition for respective time points as relative values (log_2_FC). Two genes in the original gene set were omitted from the heatmap of S phase genes since they were not expressed in all samples (see **Supplementary Table 7d**).

**Supplementary Figure 8** │ **Roles of histone acetylation and auxin biosynthesis in regulating expression of chloroplast-related genes during protoplast reprogramming.** Heatmap representing the transcriptional changes for genes implicated in chloroplast biogenesis and function in yucasin or MB-3-treated protoplasts. Genes are listed in the same order as in **Supplementary Fig. 3**.

**Supplementary Figure 9** │ **Roles of ARF7, ARF19, IAA3 and IAA18 in protoplast cell cycle reinitiation. a,** Callus formation efficiency of protoplasts isolated from WT, *arf7* and *arf19* mutants. Error bars represent standard error. n = 30 from 6 biological replicates for the WT and 15 from 3 biological replicates for the others. \*\*\**P* < 0.001 (two-tailed Student’s *t*-test or Welch’s *t*-test compared to WT). **b,** Transcriptional changes for Aux/IAA genes in Arabidopsis and their protein interaction with ARF7 and ARF19. The leftmost column shows heatmaps representing relative expression levels (log_2_FC) in protoplasts at Day 0 compared to 23 DAS leaves. The second column shows heatmaps representing relative expression levels (log_2_FC) in protoplasts at Day 2, 4, 6 and 14 compared to Day 0. Average RPM indicates the overall expression level for each gene in 23 DAS leaves or cultured protoplasts. Right column shows Aux/IAA proteins that can interact with ARF7 and/or ARF19 based on Piya *et al.*^42^. The strength of protein interaction is shown in gray and black circles.

**Supplementary Figure 10** │ **Roles of LBD16, LBD18 and LBD33 in callus formation from hypocotyl explants.** Hypocotyl explants from WT and *lbd16-1 lbd18-1 lbd33-1* seedlings were incubated on CIM for 21 days. Callus formation was assessed using more than 80 explants from 2 biological replicates for each genotype and phenotypic reproducibility was confirmed. Scale bars are 1 mm.

## Supplementary Table legends

**Supplementary Table 1** │ **Compositions of the media and reagents used in this study.**

**Supplementary Table 2** │ **Expression data for chloroplast-related genes.** Expression data of the chloroplast-related genes listed in **Supplementary Fig. 3** and **Supplementary Fig. 8** at Day 0 and Day 14 in control or MB-3 conditions (Experiment A) and at Day 0, Day 2, Day 4 and Day 6 in the control, yucasin and MB-3 conditions (Experiment B). The ’logFC’ columns show the relative expression levels (log_2_FC) at each time point compared to intact leaves (in the case of Day 0 data) or to Day 0 (in other cases).

**Supplementary Table 3** │ **Lists of the differentially expressed genes shown in** **Fig. 2c. a,** 4,679 genes that are significantly upregulated in the control condition at Day 14 compared to Day 0. **b,** 4,287 genes that are significantly upregulated in the MB-3 condition at Day 14 compared to Day 0. **c,** 1,186 genes that show significantly higher expression levels in the control condition compared to the MB-3 condition at Day 14. **d,** 395 genes that are that significantly upregulated only in the control condition but not in the MB-3 condition at Day 14 compared to Day 0 and also have significantly higher expression levels in the control condition than in the MB-3 condition at Day 14. **e,** 140 genes that are that significantly upregulated in both the control and MB-3 conditions at Day 14 from Day 0, but with significantly higher expression levels in the control condition than in the MB-3 condition at Day 14.

**Supplementary Table 4 │ The top 50 categories from GO enrichment analysis for the 535 upregulated genes.** GO categories classified as biological functions were used for the analysis.

**Supplementary Table 5 │ A list of genes involved in biosynthesis and metabolism of indole-containing compounds that show significantly stronger upregulation in the control condition compared to the MB-3 condition. a**, The genes categorized under GO terms related to indole-containing compound biosynthetic or metabolic process based on the analysis presented in **Supplementary Table 4**. The color key is the same as **Fig. 2d**. **b,** Expression data for **a**.

**Supplementary Table 6** │ **Expression data for IAA biosynthesis-related genes. a**, The expression of the genes involved in IAA biosynthesis and metabolism in intact leaves and in protoplasts at Day 0 and Day 14. Gene sets were selected based on annotations at Kyoto Encyclopedia of Genes and Genomes (KEGG) (https://www.genome.jp/kegg/kegg_ja.html) and The Arabidopsis Information Resource (TAIR) (https://www.arabidopsis.org/index.jsp). **b,** The expression of representative IAA biosynthetic genes at Day 0, Day 2, Day 4 and Day 6.

**Supplementary Table 7** │ **Expression data for cell cycle genes during the first week of protoplast culture. a,** The expression of the core cell cycle genes listed in **Fig. 4a**. **b,** The expression of the genes specifically expressed in S phase. The genes are listed in the same order as in **Supplementary Fig. 7a**. **c,** The expression of the genes specifically expressed in G2/M phase. The genes are listed in the same order as in **Supplementary Fig. 7a**. **d,** The expression of the genes specifically expressed in S phase. The genes are listed in the same order as in **Supplementary Fig. 7b**. **e,** The expression of the genes specifically expressed in G2/M phase. The genes are listed in the same order as in **Supplementary Fig. 7b**.

**Supplementary Table 8** │ **Expression data for the genes coding auxin signalling components that contribute to callus formation in tissue culture.**

**Supplementary Table 9** │ **Expression data for Aux/IAA genes in leaves and protoplasts.**

**Supplementary Table 10** │ **Primers used in this study.**

The primer sequences used for *YUC1* CDS (coding sequence) cloning and for *YUC1* RT-qPCR are from Sugawara *et al.*^70^. The primer sequences for *PP2A3* RT-qPCR are from Rymen *et al.*^69^.

## Notes

### Competing Interest Statement

The authors have declared no competing interest.

