## Supplementary Figures for "Transcriptional activation of auxin biosynthesis drives developmental reprogramming of differentiated cells"

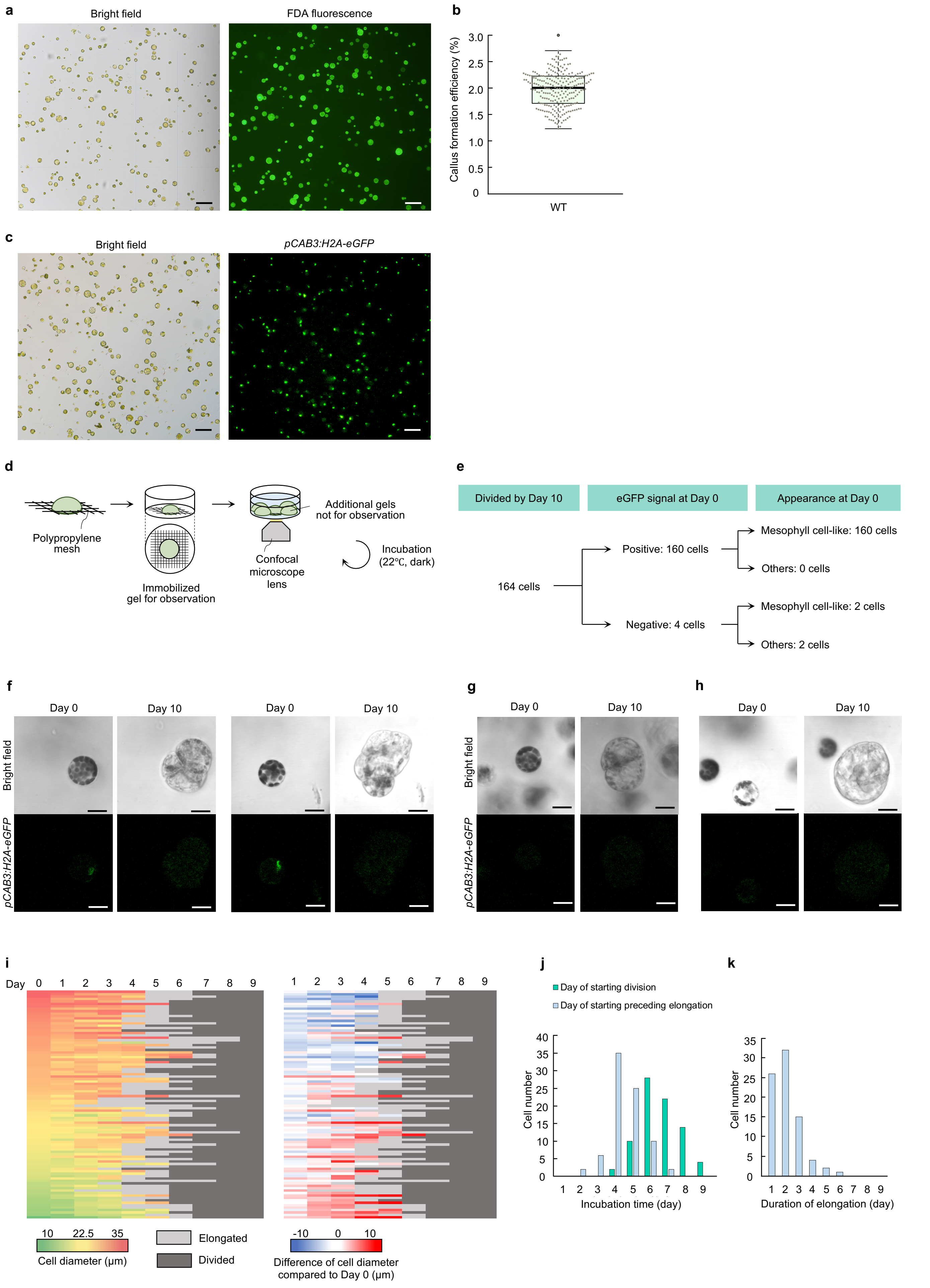


**Supplementary Figure 1**

**Supplementary Figure 1 │ Cell cycle reinitiation and callus formation described with a newly established culture system and time-lapse confocal microscopy.** **a**, Bright-field and fluorescence microscopy images of freshly isolated WT protoplasts. FDA staining indicates that 98.66 ± 0.19 % (standard error, n = 10) of freshly isolated protoplasts are viable. **b**, Callus formation efficiency of WT protoplasts. Average efficiency is 1.98 ± 0.33%. n = 240 from 48 biological replicates. Each dot shows callus formation efficiency calculated from one gel. In the box plot, the median is represented by a black line and the upper and lower quartiles are represented by the upper and lower ends of the box respectively. **c**, Bright-field and fluorescence microscopy images of protoplasts freshly isolated from plants carrying a mesophyll cell-specific marker *pCAB3:H2A-eGFP*. Quantitative analysis indicates that 94.63 ± 0.51% of freshly isolated protoplasts show distinct nuclear-localized H2A-eGFP expression (standard error, n = 15 from 2 biological replicates). **d**, Diagram showing the time-lapse confocal microscopy procedure used to track individual protoplasts. **e,** Classification of 164 protoplasts carrying *pCAB3:H2A-eGFP* that underwent cell division by Day 10 based on H2A-eGFP expression and appearance at Day 0. **f**, Two examples of *pCAB3:H2A-eGFP* protoplasts that are H2A-eGFP-positive and mesophyll cell-like at Day 0. Both protoplasts divided by Day 10. **g,** A *pCAB3:H2A-eGFP* protoplast that is H2A-eGFP-negative and mesophyll cell-like at Day 0. The protoplast divided by Day 10. **h,** A *pCAB3:H2A-eGFP* protoplast that is H2A-eGFP-negative and guard cell-like at Day 0. The protoplasts divided by Day 10. **i,** Heat maps representing cell size dynamics of protoplasts that reinitiate cell division between Day 4 and Day 9. Each row shows cell diameter (left panel) and change in cell diameter comparted to Day 0 (right panel) for individual protoplasts isolated from WT, *DR5rev:GFP* or *pVHP1:VHP1-mGFP* plants. Among 94 protoplasts tested in this experiment, 80 cells underwent cell elongation before cell division. **j,** Timing of cell elongation and cell division for 80 cells from **i**. **k,** Duration of cell elongation (right panel) for 80 cells from **i**. Scale bars are 100 µm (**a**), 100 µm (**d**) and 20 µm (**f** to **h**).

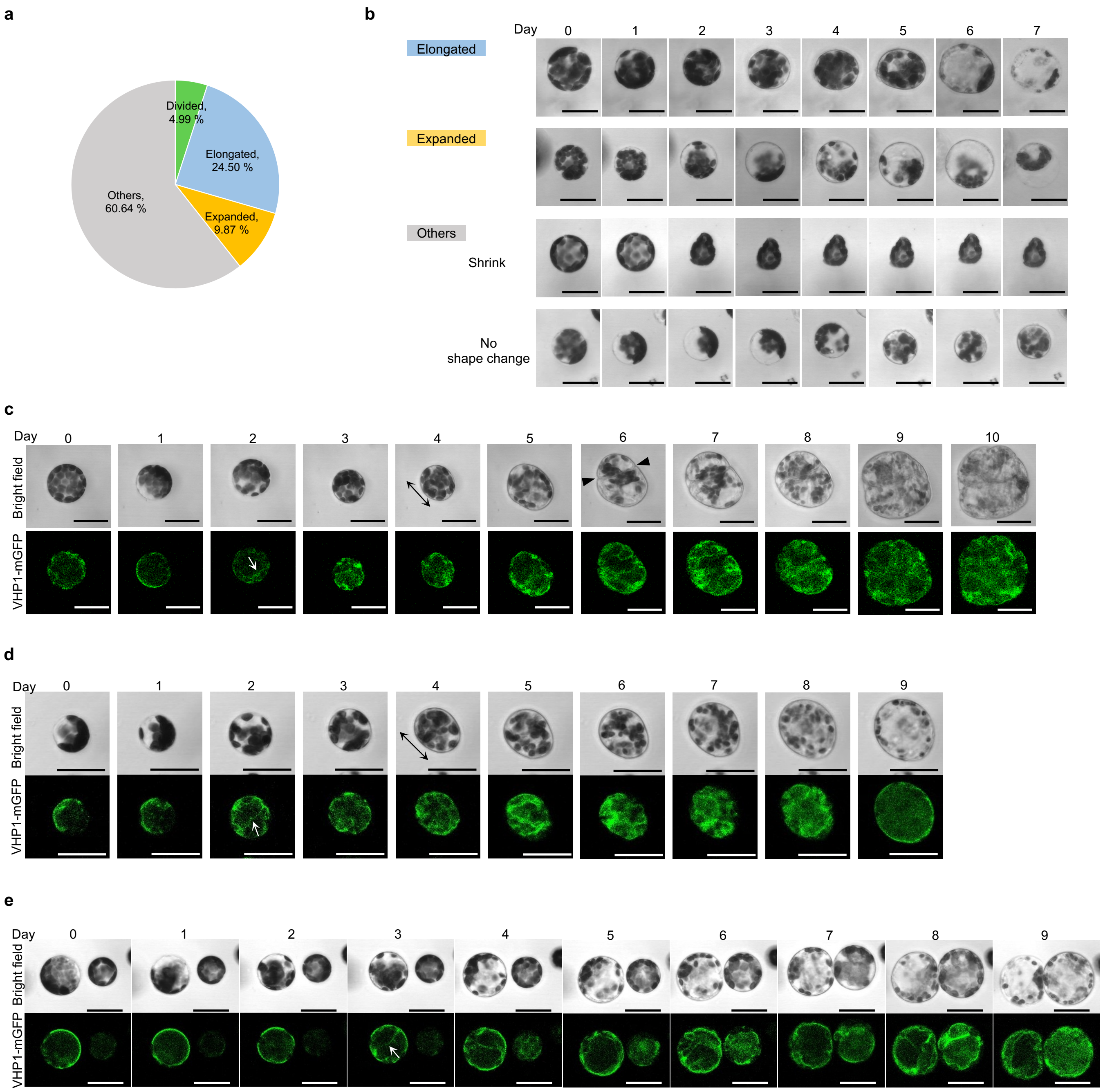


**Supplementary Figure 2 │ Time-lapse confocal microscopy images of leaf mesophyll protoplasts that did or did not undergo cell division.** **a**, Pie chart showing the percentages of protoplasts that underwent cell division (divided), those that elongated without cell division (elongated), those that expanded without cell division (expanded) and those that shrunk or displayed no shape changes (others) among 902 protoplasts isolated form *DR5rev:GFP* plants and used for time-lapse confocal microscopy. **b**, Representative time-lapse images of elongated, expanded and other protoplasts from Day 0 to Day 7. **c,** Another set of time-lapse images of a protoplast that underwent cell division. **d,** Time-lapse images of a protoplast that elongated without cell division. **e**, Time-lapse images of a protoplast that expanded without cell division. In **c** to **e**, vacuolar morphology is visualized by VHP1-mGFP. The double-headed arrow indicates the direction of cell elongation and arrowheads mark the plane of initial cell division. The white arrow highlights the initial appearance of vacuolar strand-like structures. Scale bars are 30 µm.

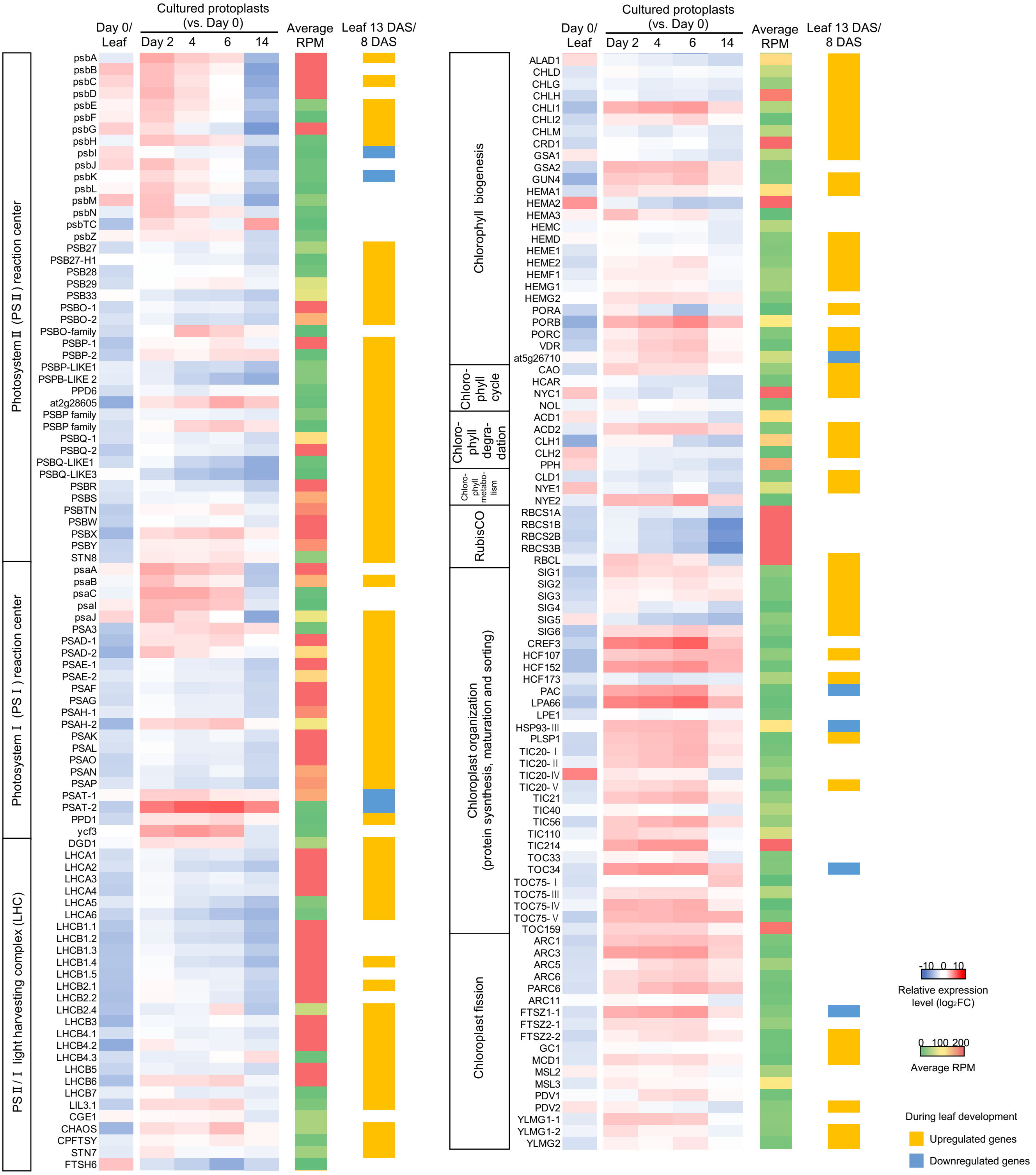


**Supplementary Figure 3 │ Transcriptional changes of chloroplast-related genes during protoplast reprogramming.** Heat map representing the transcriptional changes for genes implicated in chloroplast biogenesis and function. The leftmost column shows relative expression levels ($\log_2$FC) in protoplasts at Day 0 compared to 23 DAS leaves. The second column shows relative expression levels ($\log_2$FC) in cultured protoplasts at Day 2, 4, 6 and 14 compared to Day 0. Average reads per million (RPM) indicates the overall expression level for each gene. The right column shows genes significantly upregulated (orange) or downregulated (blue) during leaf development based on Andriankaja et al. (2012). Gene sets were selected based on annotations at The Arabidopsis Information Resource (TAIR) (https://www.arabidopsis.org/index.jsp).

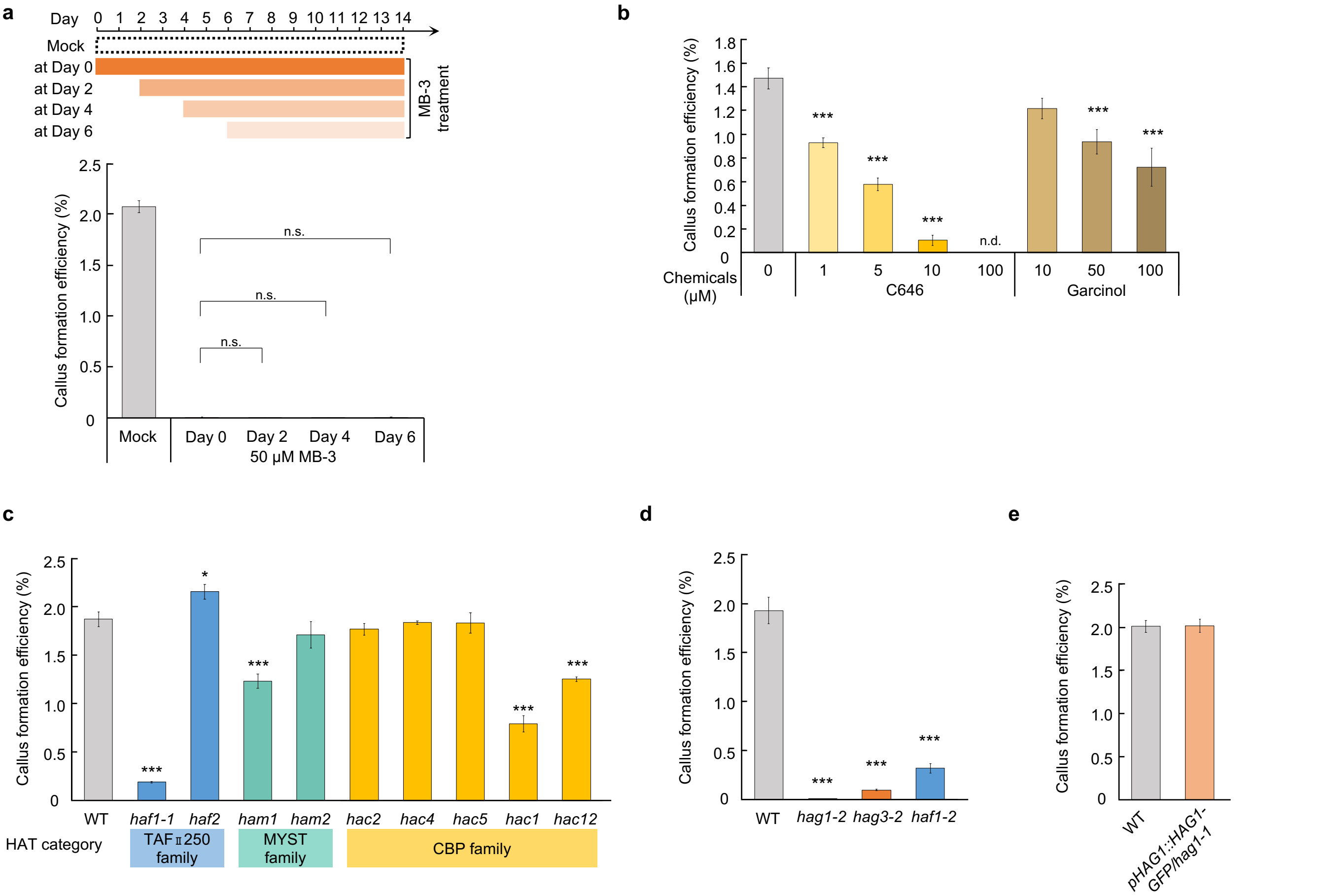


**Supplementary Figure 4 │ Roles of histone acetyltransferases in protoplast cell cycle reinitiation. a**, Callus formation efficiency of WT protoplasts treated with 50 µM MB-3 at different time points. Diagram shows the timing of MB-3 treatment. Error bars represent standard error. n = 15 from 3 biological replicates. n.s. not significant (two-tailed Student's *t*-test or Welch's *t*-test compared to MB-3 treatment at Day 0). **b**, Callus formation efficiency of WT protoplasts treated with C646 or garcinol. Error bars represent standard error. n = 30 from 6 biological replicates for the WT control and n = 15 from 3 biological replicates for all others. ***$P < 0.001$ (two-tailed Student's *t*-test or Welch's *t*-test compared to WT control). n.d. not determined. **c**, Callus formation efficiency of protoplasts isolated from WT and HAT mutants. Error bars represent standard error. n = 45 from 9 biological replicates for the WT and n = 10 or 15 from 2 or 3 biological replicates for mutants. *$P < 0.05$, *** $P < 0.001$ (two-tailed Student's *t*-test or Welch's *t*-test compared to WT). **d**, Callus formation efficiency of WT, *hag1-2*, *hag3-2* and *haf1-2* protoplasts. Error bars represent standard error. n = 25 from 5 biological replicates for the WT and n = 15 from 3 biological replicates for all others. *** $P < 0.001$ (two-tailed Welch's *t*-test compared to WT). **e**, Callus formation efficiency of WT and *pHAG1:HAG1-GFP/hag1-1* protoplasts. Error bars represent standard error. n = 15 from 3 biological replicates. No statistical difference was detected (two-tailed Student's *t*-test compared to WT).

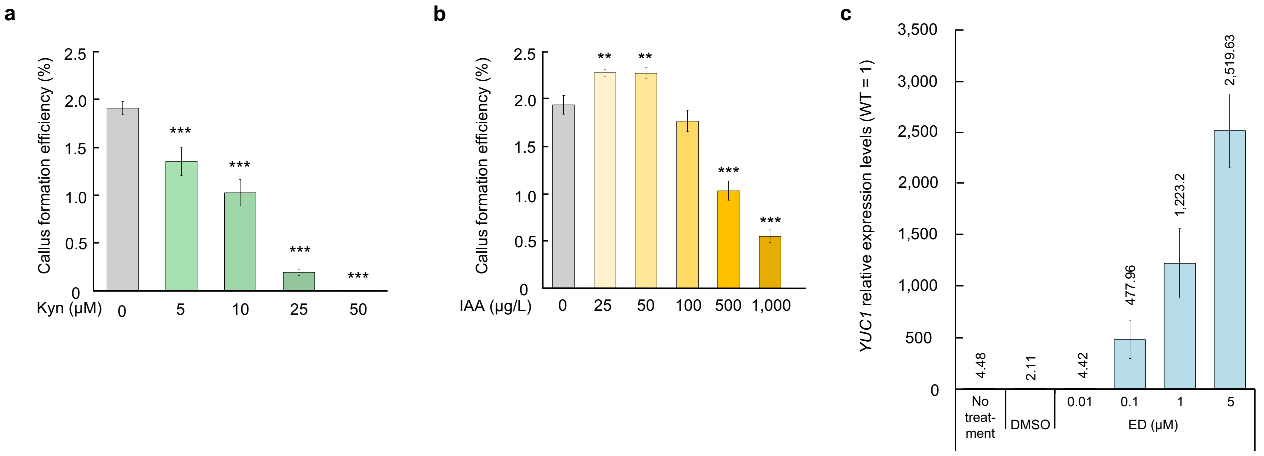


**Supplementary Figure 5│ Roles of auxin biosynthesis in protoplast cell cycle reinitiation. a**, Callus formation efficiency of Kyn-treated WT protoplasts. Error bars represent standard error. n = 30 from 6 biological replicates for the WT control and n = 15 to 25 from 3 to 5 biological replicates. ***$P < 0.001$ (two-tailed Student's *t*-test or Welch's *t*-test compared to WT control). **b**, Callus formation efficiency of WT protoplasts incubated with PCIM supplemented with 0 to 1,000 μg/L IAA. Error bars represent standard error. n = 10 to15 from 2 to 3 biological replicates. **$P < 0.01$, ***$P < 0.001$ (two-tailed Student's *t*-test or Welch's *t*-test compared to WT control). **c**, β-estradiol-inducible expression of *YUC1* in *XVE-YUC1* plants. WT and *XVE-YUC1* seeds were sown in liquid half-strength MS medium and incubated at 22℃ under light with rotation on a Shake-LR (TAITEC) for 10 days. The plants were then treated with different concentrations of ED or the equivalent volumes of DMSO and grown for another 24 hours. The expression levels of *YUC1* are normalized by those of the internal control *PP2A3* and shown as relative values compared to WT given the same treatments. The *YUC1* expression in WT remained constant throughout the treatments. The numbers above each bar are average expression levels. Error bars represent standard error (n = 6).

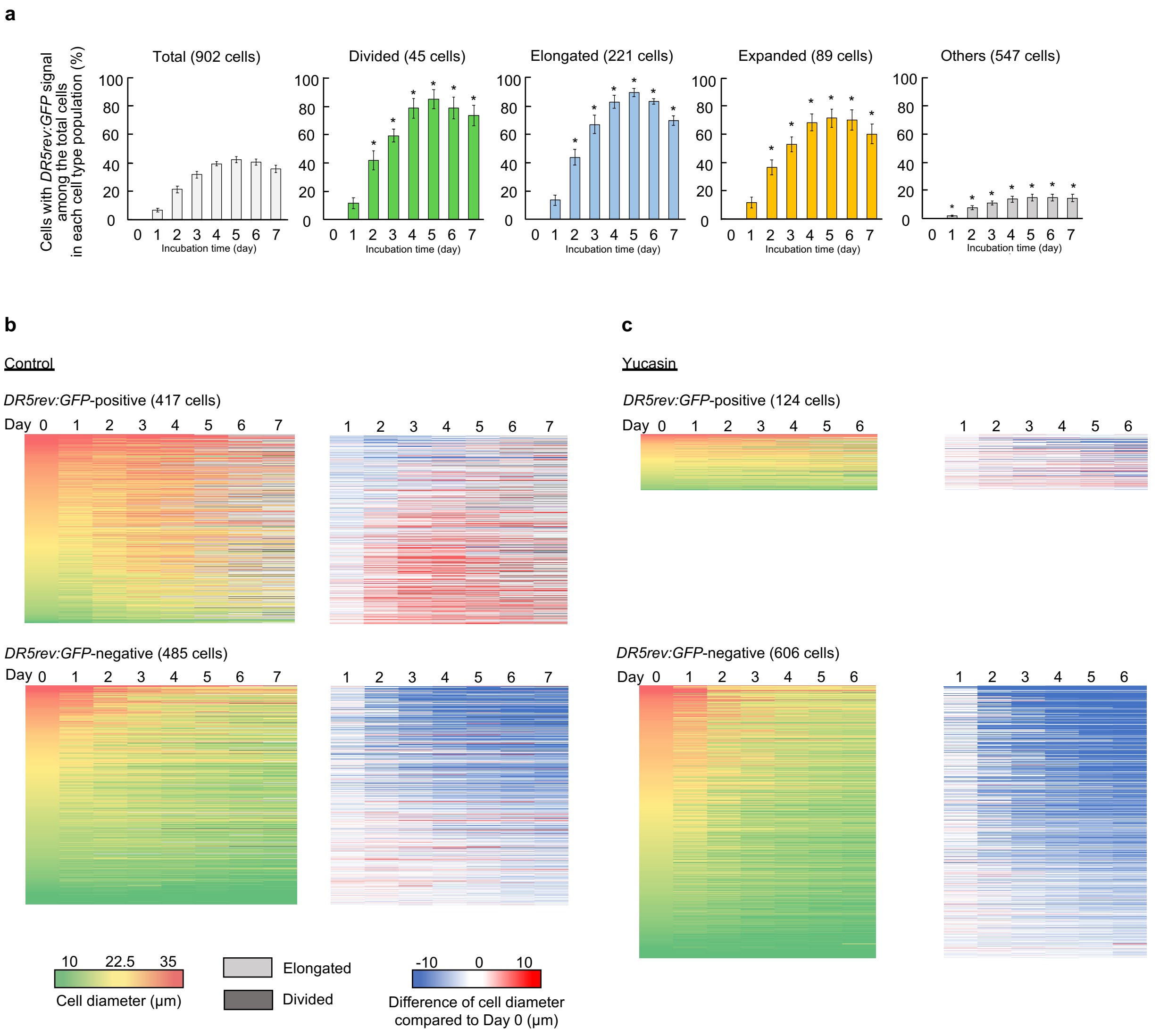


**Supplementary Figure 6 │ Correlation between auxin response and the morphological changes during reprogramming of protoplasts. a**, Frequency of the *DR5rev:GFP*-expressing (positive) protoplasts among 902 total cells, 45 divided cells, 221 elongated cells, 89 expanded cells and 547 other cells. *DR5rev:GFP* protoplasts were incubated in the control condition. **P* < 0.05 (two-tailed Student's *t*-test or Welch's *t*-test compared to 'Total' at the same time point). **b**, Heat maps representing cell size dynamics of 417 *DR5rev:GFP*-positive and 485 *DR5rev:GFP*-negative protoplasts in the control condition. **c,** Heat maps representing cell size dynamics of 124 *DR5rev:GFP*-positive and 606 *DR5rev:GFP*-negative protoplasts in the 50 μM yucasin condition. In **b** and **c**, each row shows cell diameter (left panel) and change in cell diameter comparted to Day 0 (right panel) for individual *DR5rev:GFP* protoplasts.

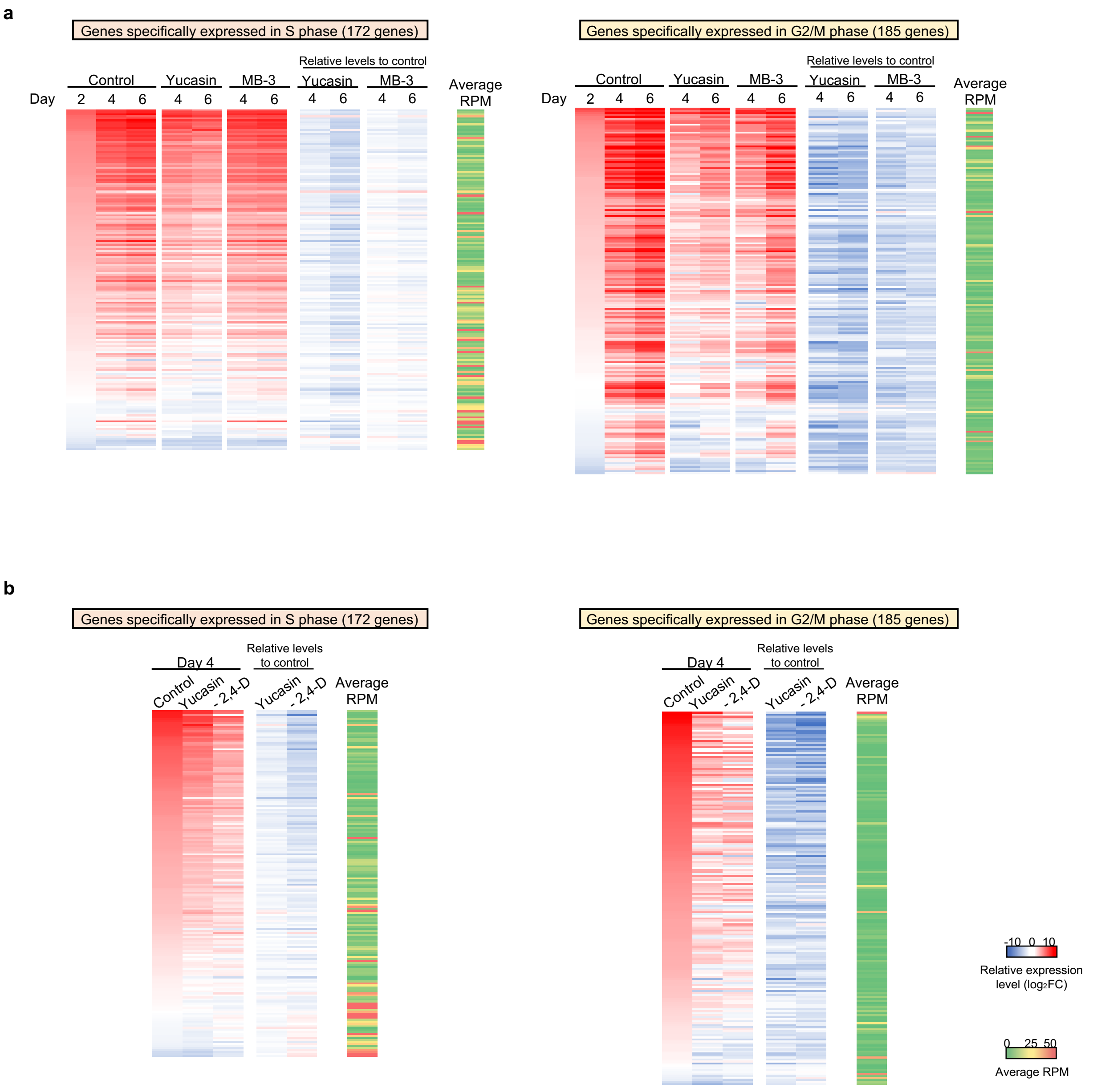


**Supplementary Figure 7 │ Roles of histone acetylation, auxin biosynthesis and exogenous 2,4-D in regulating expression of cell cycle genes during protoplast reprogramming. a**, Heatmap representing the transcriptional changes for 172 genes specifically expressed at S phase and 185 genes specifically expressed at G2/M phase. The left three columns show expression levels in the control, 50 µM yucasin, and 50 µM MB-3 condition as values normalized ($log_2$FC) to Day 0. The 'Relative levels to control' columns show the normalized expression levels in the yucasin or MB-3 condition compared to the control condition for respective time points as relative values ($log_2$FC). Average RPM indicates the overall expression level for each gene. Gene sets are taken from Kobayashi *et al.*[29]. Two genes in the original gene set were omitted from the heatmap of S phase genes since they were not expressed in all samples (see **Supplementary Table 7b**). **b,** Heatmap representing the transcriptional changes for the same gene sets as in **a**. The left three columns show expression levels at Day 4 in the control, 50 µM yucasin, and 2,4-D-omitted (named '- 2,4-D') condition as values normalized ($log_2$FC) to Day 0. The 'Relative levels to control' columns show the normalized expression levels in yucasin or - 2,4-D condition compared to the control condition for respective time points as relative values ($log_2$FC). Two genes in the original gene set were omitted from the heatmap of S phase genes since they were not expressed in all samples (see **Supplementary Table 7d**).

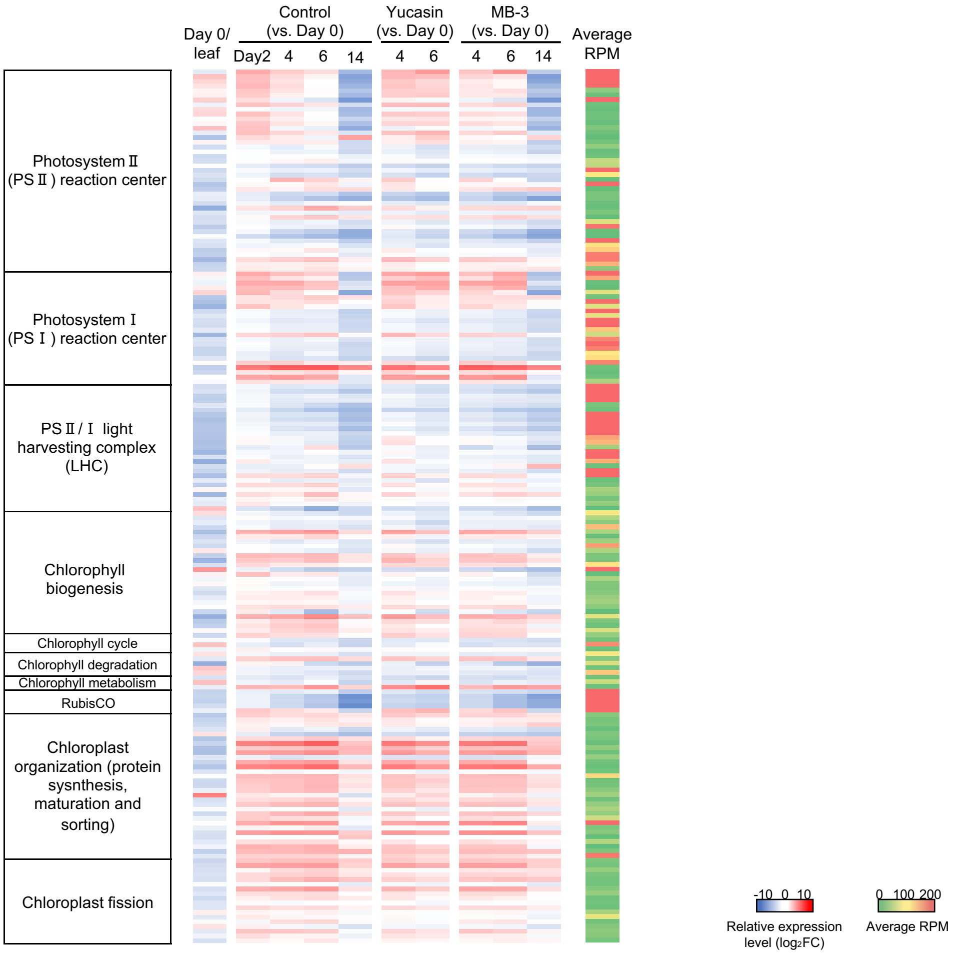


**Supplementary Figure 8 │ Roles of histone acetylation and auxin biosynthesis in regulating expression of chloroplast-related genes during protoplast reprogramming.** Heatmap representing the transcriptional changes for genes implicated in chloroplast biogenesis and function in yucasin or MB-3-treated protoplasts. Genes are listed in the same order as in **Supplementary Fig. 3**.

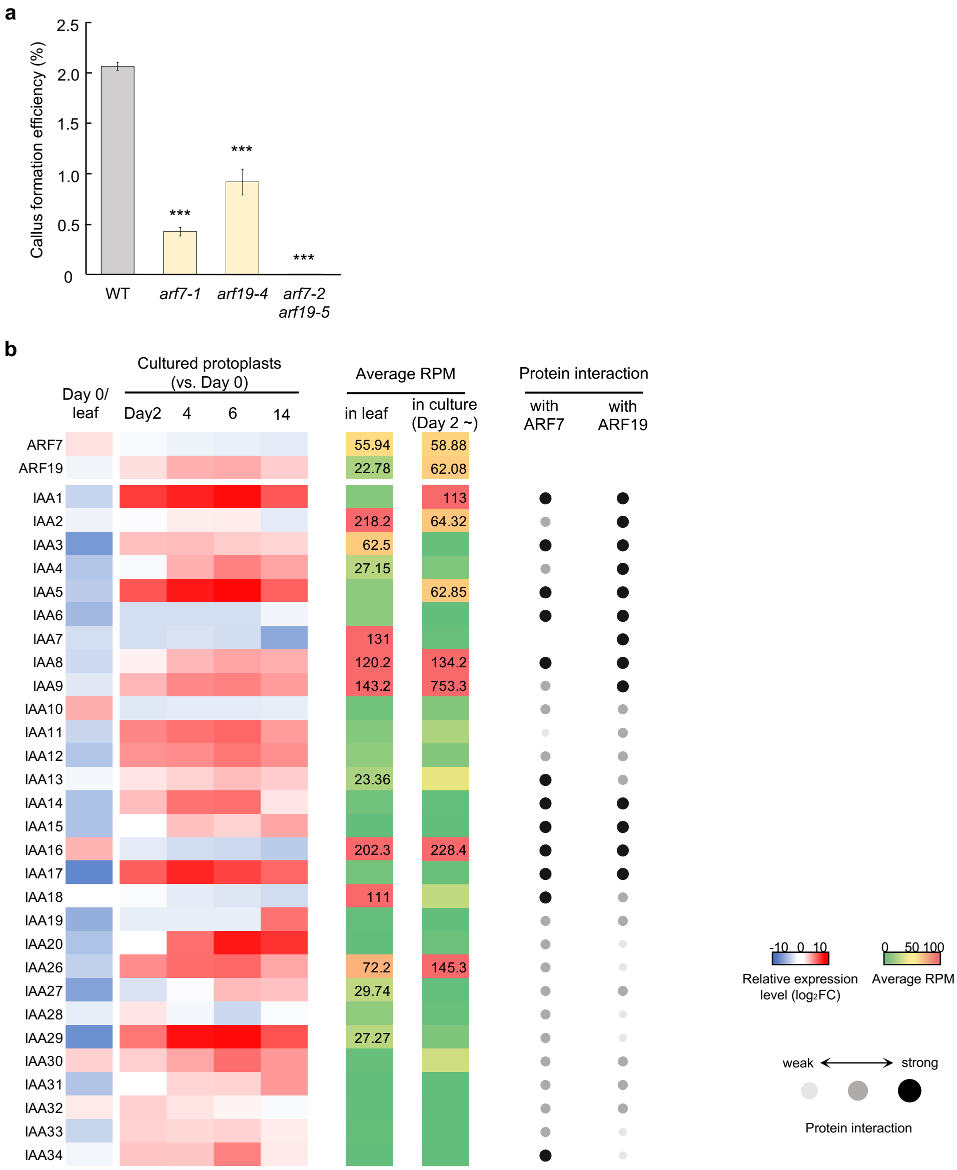


**Supplementary Figure 9 │ Roles of ARF7, ARF19, IAA3 and IAA18 in protoplast cell cycle reinitiation. a,** Callus formation efficiency of protoplasts isolated from WT, *arf7* and *arf19* mutants. Error bars represent standard error. n = 30 from 6 biological replicates for the WT and 15 from 3 biological replicates for the others. ***$P < 0.001$ (two-tailed Student's *t*-test or Welch's *t*-test compared to WT). **b,** Transcriptional changes for Aux/IAA genes in Arabidopsis and their protein interaction with ARF7 and ARF19. The leftmost column shows heatmaps representing relative expression levels ($\log_2$FC) in protoplasts at Day 0 compared to 23 DAS leaves. The second column shows heatmaps representing relative expression levels ($\log_2$FC) in protoplasts at Day 2, 4, 6 and 14 compared to Day 0. Average RPM indicates the overall expression level for each gene in 23 DAS leaves or cultured protoplasts. Right column shows Aux/IAA proteins that can interact with ARF7 and/or ARF19 based on Piya *et al.*[42]. The strength of protein interaction is shown in gray and black circles.

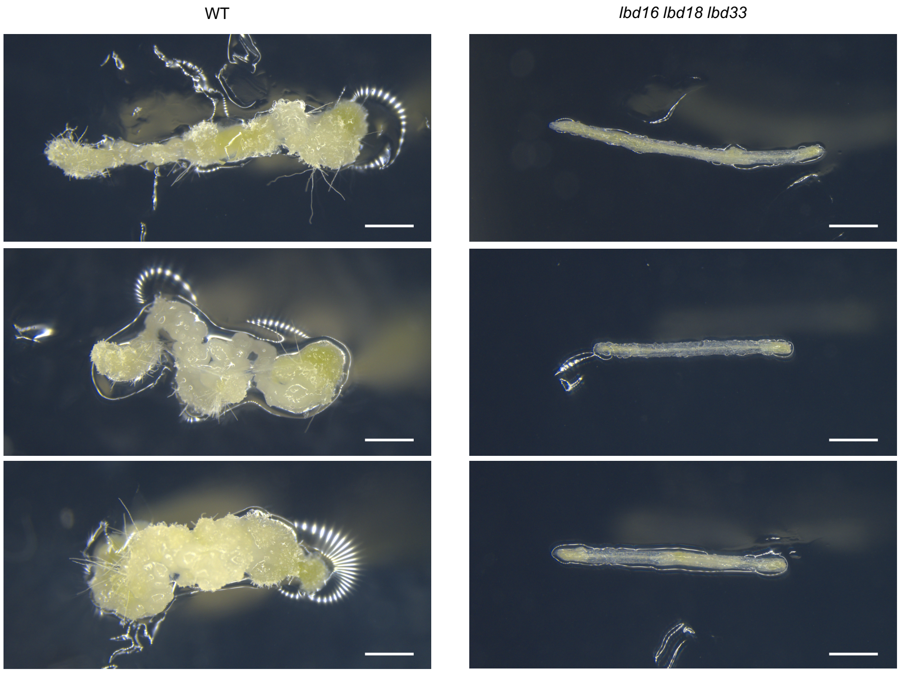


**Supplementary Figure 10 │ Roles of LBD16, LBD18 and LBD33 in callus formation from hypocotyl explants.** Hypocotyl explants from WT and *lbd16-1 lbd18-1 lbd33-1* seedlings were incubated on CIM for 21 days. Callus formation was assessed using more than 80 explants from 2 biological replicates for each genotype and phenotypic reproducibility was confirmed. Scale bars are 1 mm.
